# Furry is required for cell movements during gastrulation and functionally interacts with NDR1

**DOI:** 10.1101/2020.05.08.083980

**Authors:** Ailen S. Cervino, Bruno Moretti, Carsten Stuckenholz, Hernán E. Grecco, Lance A. Davidson, M. Cecilia Cirio

## Abstract

Gastrulation is a key event in animal embryogenesis during which the germ layers precursors are rearranged and the embryonic axes are established. Cell polarization is essential during gastrulation driving asymmetric cell division, cell movements and cell shape changes. Furry (Fry) gene encodes an evolutionarily conserved protein with a wide variety of cellular functions mostly related to cell polarization and morphogenesis in invertebrates. However, little is known about its function in vertebrate development. Here we show that in *Xenopus*, Fry participates in the regulation of morphogenetic processes during gastrulation. Using morpholino knock-down, we demonstrate a role of Fry in blastopore closure and dorsal axis elongation. Loss of Fry function drastically affects the movement and morphological polarization of cells during gastrulation, in addition to dorsal mesoderm convergent extension, responsible for head-to-tail elongation. Finally, we demonstrate a functional interaction between Fry and NDR1 kinase, providing evidence of an evolutionarily conserved complex required for morphogenesis.

## Introduction

Gastrulation is a crucial time in animal development during which major cell and tissue movements shape the basic body plan (Keller et al., 2003; Leptin, 2005). The morphogenetic movements of gastrulation rearrange the three germ layers precursors positioning the mesodermal cells between the outer ectodermal and the inner endodermal cells, and shape the head-to-tail body axis. In order to break the initial “egg shape” of the embryo, cells need to polarize in a precise and coordinated manner. Cell polarity controls orientated cell division, cell shape changes, as well as cell movement and it determines the mechanical properties of the tissue, e.g. extracellular matrix (ECM) organization during gastrulation and numerous other morphogenetic events (Keller, 2002; Keller and Shook, 2008; Keller et al., 2003). The frog *Xenopus laevis* has been widely used as a model of cell polarization, migration, and morphogenesis due to its unique experimental advantages, such as the large size of the embryo and its cells, allowing extensive manipulation and high resolution live microscopy of explant cultures (Keller and Shook, 2008; Shih and Keller, 1992a).

At the beginning of *Xenopus* gastrulation, the presumptive anterior mesoderm cells located at the dorsal marginal zone (DMZ) roll inward at the midline of the blastopore lip in a process called involution. Involution follows bottle cell contraction and spreads laterally and ventrally leading to the formation of the blastopore, a ring of involuting cells that encircles the yolky vegetal endoderm cells. As involution proceeds, the blastopore progressively decreases in diameter, defining the posterior of the embryo, and closes at the end of gastrulation (Keller et al., 2003). Simultaneously, on the dorsal side of the embryo, the axial and paraxial mesoderm cells undergo convergent extension which elongates the antero-posterior axis and aids blastopore closure. During convergent extension, mesodermal cells need to polarize and intercalate with each other along the mediolateral axis, narrowing and extending the dorsal midline (Keller et al., 2000; Wallingford et al., 2002).

The Furry (Fry) gene encodes a large protein that is conserved evolutionarily from yeast to humans. Fry protein is composed of an N-terminal Furry domain (FD) with HEAT/Armadillo repeats and five less well conserved regions without any recognizable functional domains. Additionally, there are two leucine zipper motifs and a coiled-coil motif at the C-terminus in vertebrates only (Nagai and Mizuno, 2014). The phenotypes associated with the loss-of-function of Fry orthologs in invertebrates, including *Drosophila* Fry, *C.elegans* Sax-2, *S. pombe* Mor2p and *S. cerevisiae* Tao3p, suggest that this protein is implicated in the control of cell division, transcriptional asymmetry, cell polarization and morphogenesis (Cong et al., 2001; Du and Novick, 2002; Gallegos and Bargmann, 2004; He et al., 2005; Hirata et al., 2002; Horne-Badovinac et al., 2012; Nelson et al., 2003; Zallen et al., 2000). In mammalian cells, Fry was found in association with microtubules regulating chromosome alignment and bipolar spindle formation in mitosis (Chiba et al., 2009; Ikeda et al., 2012; Nagai et al., 2013). Many of Fry functions in vertebrates and invertebrates are related to its function as essential scaffolding factor for large protein complexes that contain NDR (<u>n</u>uclear <u>D</u>bf-2-<u>r</u>elated) protein kinases. Genetic and physical interactions between Fry and NDR1 have been observed across a broad group of eukaryotes where Fry protein acts to modulate NDR1 phosphorylation and kinase activity (Chiba et al., 2009; Gallegos and Bargmann, 2004; He et al., 2005; Hergovich et al., 2006; Xiong et al., 2018). In *Xenopus*, the function of Stk38, ortholog of NDR1, nor its physical and functional interaction with Fry during development has been studied.

Fry’s role in vertebrate development has only been studied in *Xenopus*. It was found as a maternally expressed gene (Goto et al., 2010). In the early gastrula embryo, *fry* transcripts are present in the dorsal and ventral tissues and later in the mesoderm and ectoderm derivatives (Espiritu et al., 2018; Goto et al., 2010). Fry function has been associated with the regulation of microRNA expression in chordamesoderm development and the specification of the pronephric kidney (Espiritu et al., 2018; Goto et al., 2010). In this study we investigate the participation of Fry in morphogenetic processes that occur during gastrulation. We first described its expression during gastrulation in *Xenopus* embryos and using morpholino knock-down, we show that Fry is required for blastopore closure and axis elongation. At the cellular level, loss of Fry function drastically affects the movement and morphological polarization of mesodermal cells during gastrulation. Moreover, convergent extension of the dorsal mesoderm, which is one of the major tissue movements that contribute to blastopore closure and extension of the body axis, is impaired in *fry*-depleted embryos. We further show a functional interaction between Fry and NDR1 kinase, providing evidence of an evolutionarily conserved requirement of these proteins in morphogenesis.

## Results

### Fry depletion causes axis elongation defects without mesoderm induction impairment

We and others have previously determined that *fry* is expressed in the involuting mesoderm of early gastrula, becoming restricted to dorsal tissues and lateral plate mesoderm of neurula, and remaining in somites, notochord, heart, eye, brain and pronephric kidney through tailbud stages (Espiritu et al., 2018; Goto et al., 2010). We investigated in more detail its expression pattern before and during gastrulation and found that *fry* transcripts are present almost exclusively in the animal half of the blastula and the expression domain expands to the marginal zone in early gastrula (Fig. S1A-B). By late gastrula, *fry* transcripts, are present in the axial mesoderm and the deep layer of the ectoderm becoming restricted to the notochord, paraxial and lateral mesoderm in neurula (Fig. S1C-D). We went further and decided to look at Fry cellular localization in DMZ explants prepared from embryos injected with *fry-GFP* mRNA at a dose that did not cause an axis phenotype. Since we were not able to detect the endogenous protein with available antibodies, we use a Fry-GFP fusion protein (Goto et al., 2010) and found it mainly present in the cytoplasm and membrane of dorsal mesodermal cells during gastrulation (Fig. S1E). The morphogenetic defects associated with the loss of Fry function in invertebrates and our results of transcript and protein localization in the *Xenopus* embryo, led us to investigate a role of Fry in morphogenetic movements at gastrulation.

We performed loss-of-function experiments by knocking down *fry* translation using a previously-validated antisense morpholino oligonucleotide (*fry-*MO) (Goto et al., 2010). As reported, injection of *fry-*MO into both dorsal blastomeres of 4-cell embryos resulted in shortened anterior-posterior axis and reduced anterior head structures at the tailbud stage (Fig. 1A-D; Goto et al., 2010). Here we show that this effect was dose dependent and we classified embryos as “Shortened axis” or “Shortened axis + Head-less” as a more severe phenotype when the cement gland and the optic and otic vesicles were absent (Fig. 1C,D,G). Additionally, we could rescue this *fry*-depletion phenotype by coinjection of *FD+LZ* mRNA that lacks the morpholino target site (Fig. 1E,G). FD+LZ is a short chimeric version of Fry that only possess the N-terminal FD and C-terminal LZ domains (Goto et al., 2010). Similar to our previous report on Fry function in the kidney (Espiritu et al., 2018), our results here indicate that together, the FD and LZ domains of Fry, are sufficient to rescue the loss-of-function short-axis phenotype. Intriguingly, dorsal overexpression of *FD+LZ* mRNA alone caused axis elongation impairment and phenocopies Fry loss-of-function phenotype indicating that normal Fry function is required for axis development (Fig. 1F,G).

**Figure 1.**
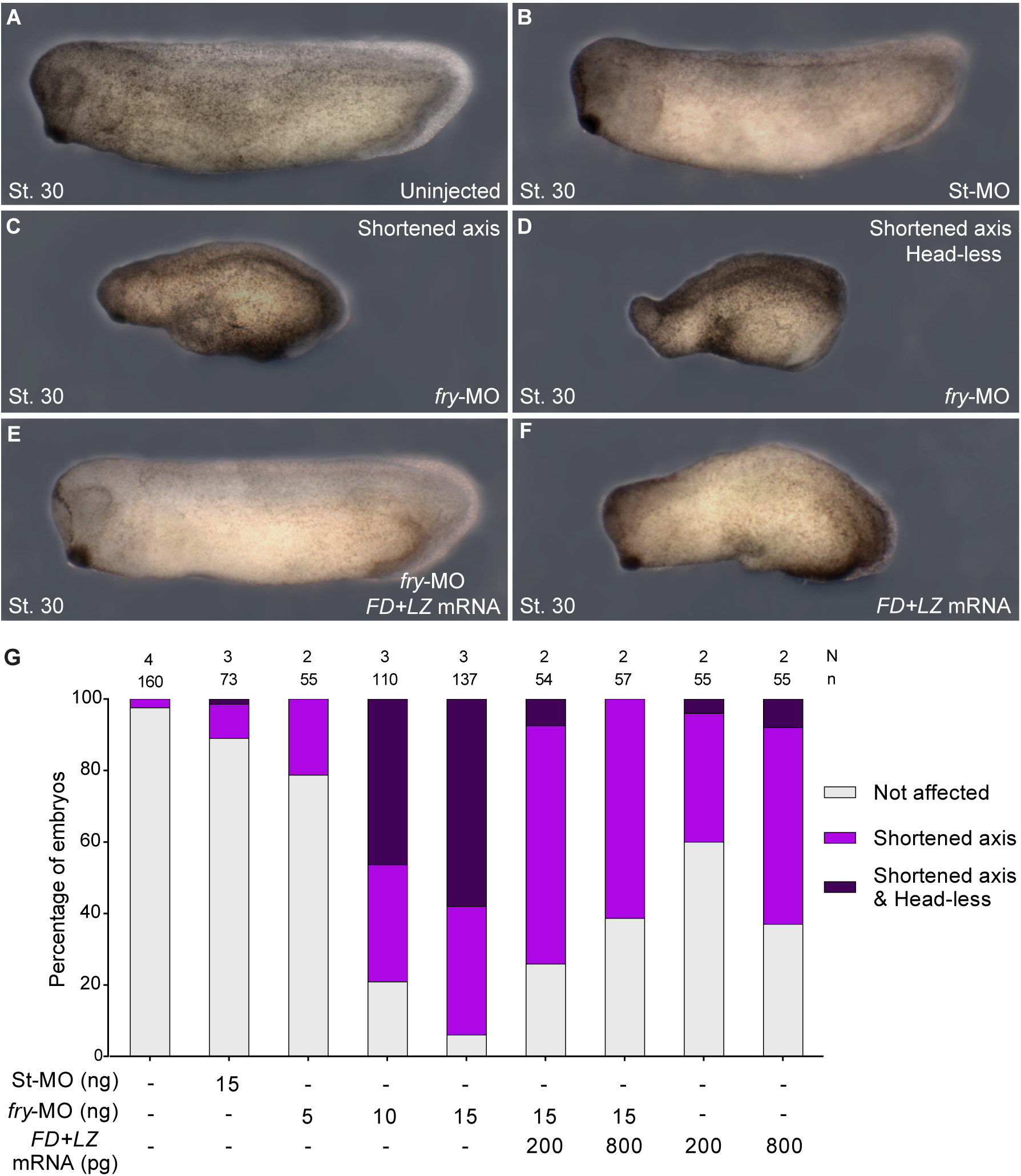
Fry loss-of-function, phenotype and rescue experiments. 4-cell stage *Xenopus* embryos were injected into both dorsal blastomeres as indicated and fixed at stage 30 (St. 30). (**A**) Uninjected embryo. (**B**) Standard Control morpholino (St-MO) (15 ng) injected embryo. (**C** and **D**) *fry*-MO (15 ng) injected embryos exhibiting “Shortened axis” or “Shortened axis & Head-less” phenotypes, respectively. (**E**) Rescue experiment; *fry*-MO (15 ng) + *FD+LZ* mRNA (800 pg) coinjected embryo. (**F**) *FD+LZ* mRNA (800 pg) injected embryo. (**G**) Quantitation of the percentage of embryos showing the different phenotypes: “Not affected”, “Shortened axis” or “Shortened axis & Head-less”. N: number of independent experiments, n: number of embryos. Data on graph is presented as mean. Representative embryos are shown.

Since Fry indirectly regulates chordamesodermal gene expression (Goto et al., 2010), we asked whether abnormal axis development was caused by defective mesoderm induction and patterning. The expression of the pan-mesodermal marker *brachyury (xbra)* (Smith et al., 1991) was present in the absence of Fry at gastrula stages, indicating that mesoderm induction was not impaired (Fig. 2A,B). Additionally, in early gastrula, expression patterns of *xbra* and the early chordamesoderm gene *notochord homeobox, not* (Von Dassow et al., 1993) appeared similar in uninjected and *fry*-MO injected embryos (Fig. 2A,B,E,F). By late gastrula stages their expression domains were abnormal. However, these shifts are consistent with defects in gastrulation movements (Fig. 2C,D,G,H). In agreement with this interpretation, the expression domains of the chordamesoderm gene *chordin* and the paraxial mesoderm gene *myoD* were not affected in the absence of Fry at late tailbud stage (Fig. S2A-D). Moreover, the notochord and somites of *fry*-morphants were similarly detected by immunostaining with MZ15 and 12/101 antibodies, indicating that differentiation of these tissues is not impaired (Fig. S2E-H). Gene expression, immunostaining and histological sections of *fry* knock-down embryos are consistent with defects in axial tissue morphogenesis (Figs 2, S2). Therefore, we decided to test the role of Fry in gastrulation movements.

**Figure 2.**
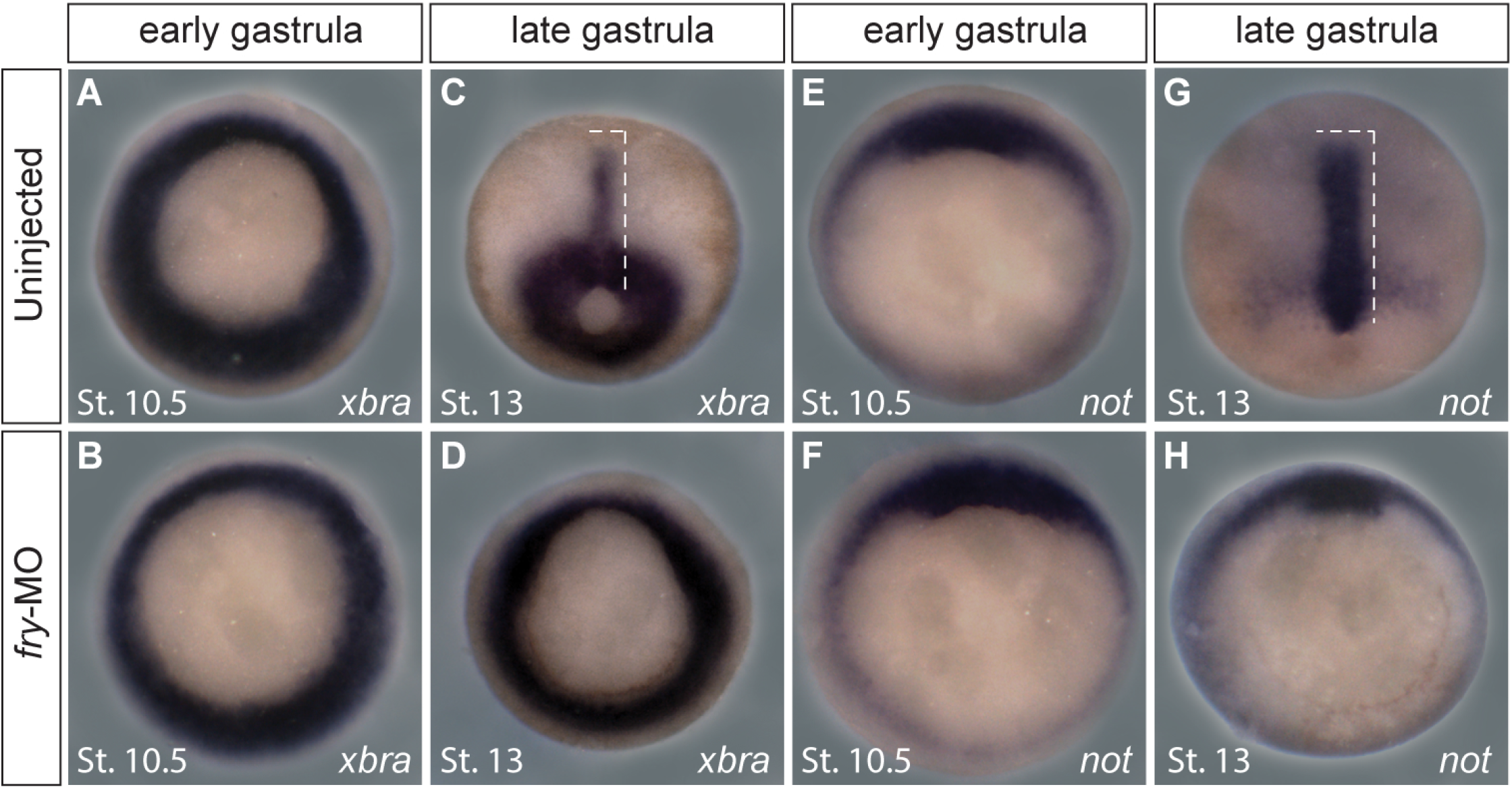
Fry depletion affects normal gastrulation movements. (**A**-**D**) *In situ* hybridization for the pan-mesoderm gene *brachyury (xbra)* in *Xenopus* embryos. (**A, B**) Early gastrula stage embryos (St. 10.5). (**A**) Uninjected embryo (N = 3; n = 47). (**B**) *fry*-MO (15 ng) injected embryo (N = 3; n = 53). (**C, D**) Late gastrula stage embryos (St. 13). (**C**) Uninjected embryo (N = 2; n = 44). (**D**) *fry*-MO (15 ng) injected embryo (N = 2; n = 52). (**E**-**H**) *In situ* hybridization for the chordamesoderm gene *notochord homeobox (not)* in *Xenopus* embryos. (**E, F**) Early gastrula stage embryos (St. 10.5). (**E**) Uninjected embryo (N = 2; n = 52). (**F**) *fry*-MO (15 ng) injected embryo (N = 2; n = 47). (**G**, **H**) Late gastrula stage embryos (St. 13). (**G**) Uninjected embryo (N = 3; n = 39). (**H**) *fry*-MO (15 ng) injected embryo (N = 3; n = 43). The stage of injected embryos was established based on the stage of uninjected littermates. Dashed lines indicate dorsal mesoderm elongation. Representative embryos are shown.

### Furry is necessary for normal blastopore closure

At the start of gastrulation, the prospective dorsal mesoderm located in the marginal zone rolls inward over the dorsal blastopore lip (DBL) in a process termed involution. As gastrulation proceeds, the blastopore closes as post-involution dorsal mesoderm extends along the anterior-posterior axis and narrows along the mediolateral axis driving axis elongation (Shih and Keller, 1992b). Thus, we measured blastopore area on fixed embryos at different stages to assess blastopore closure progression from the initial step of blastopore formation at the early gastrula, until its closure at the early neurula (Fig. 3A). Although the blastopore remained open at the late gastrula stage, most of *fry*-depleted embryos achieved blastopore closure during neurulation suggesting a delay in dorsal morphogenesis (Fig. 3A). Consistent with our initial rescue experiments observed by tailbud (Fig. 1), the blastopore closure defect induced by *fry* knock-down was partially rescued at the late gastrula stage by coinjection of *FD+LZ* mRNA (Fig. 3A). To further characterize the dynamics of blastopore closure, we acquired time-lapse sequences of gastrulating embryos from the onset of gastrulation until blastopore closure (Movie S1). We noticed that blastopore formation, which is initially restricted to a small region on the dorsal side of the uninjected embryo (t = 105 min), was retarded and laterally expanded in *fry*-morphants (t = 120 min) (yellow dotted arrows, Fig. 3B,C). Additionally, by late gastrula stages, as opposed to the characteristic dorsal-dominated eccentric closure of the blastopore toward the ventral side, *fry*-depleted embryos close their blastopore concentrically (see Movie S1 at t = 594 min and t = 480 min, Fig. 3B,C). Hence, Fry depletion in dorsal tissues alters blastopore formation, the dynamics of blastopore closure, and the position of the closing blastopore.

**Figure 3.**
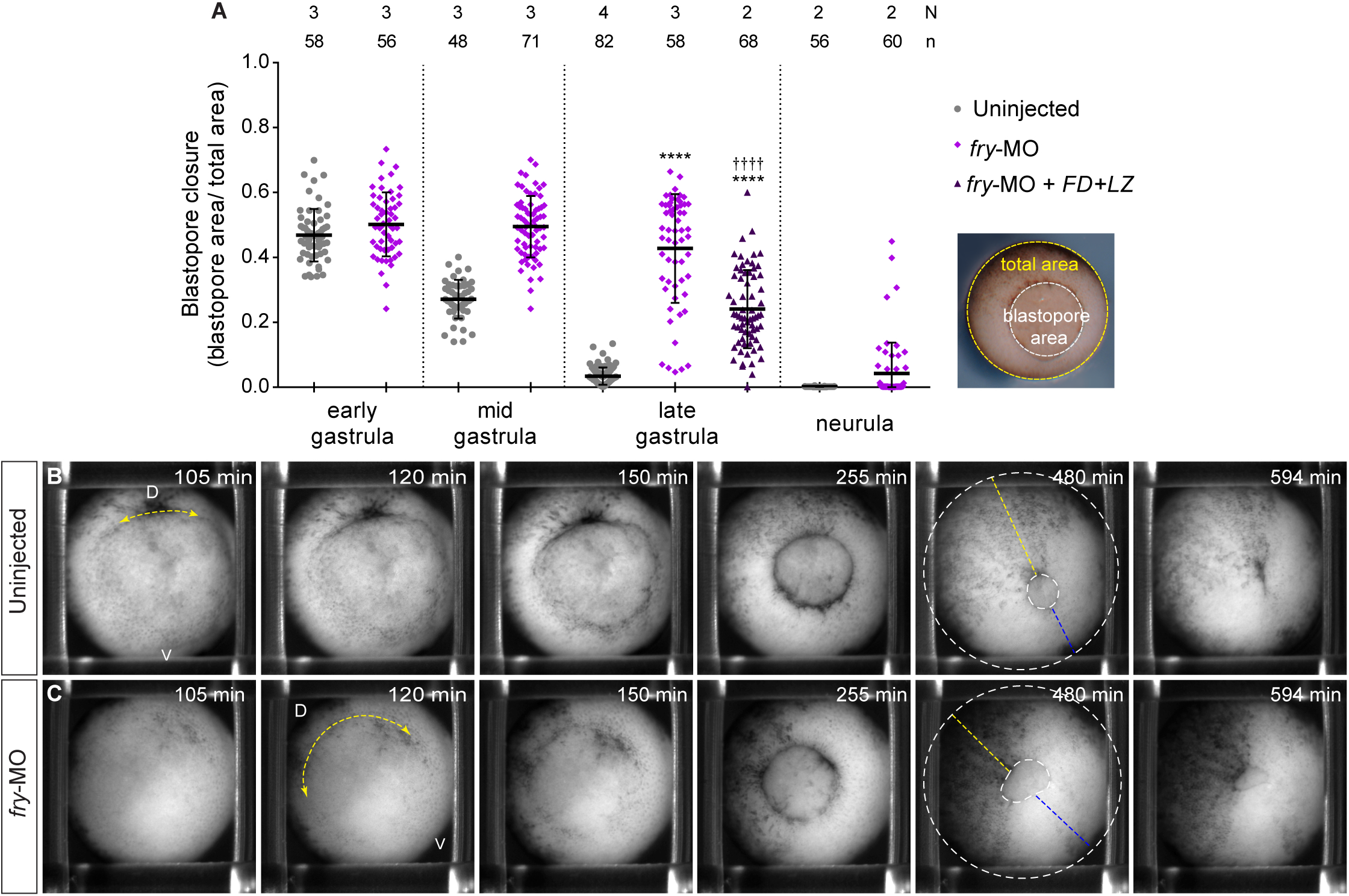
Fry is required for normal blastopore closure. (**A**) Left: Quantification of the blastopore closure measurements on uninjected embryos, *fry*-MO (15 ng) injected embryos and *fry*-MO (15 ng) + *FD+LZ* mRNA (800 pg) coinjected embryos at early gastrula stage (St. 10.5), mid gastrula stage (St. 12), late gastrula stage (St. 13) and neurula stage (St. 15). Right: Scheme of the metrics used: ratio of the blastopore area (white) over the area of the vegetal hemisphere of the embryo (total area, yellow). 0 = blastopore closed. N: number of independent experiments, n: number of embryos. The stage of injected embryos was established based on the stage of uninjected littermates. Means and standard deviation are indicated. Each point represents a single embryo. * and † indicate statistically significant differences relative to uninjected and *fry*-MO injected group, respectively. Kruskal-Wallis test and Dunn’s multiple comparisons test (**** *p*<0.0001). (**B**, **C**) Still frames from time-lapse movies (see Movie S1) of gastrulating embryos at the indicated time-point (t) (vegetal view). Embryos were mounted at late blastula (St. 9) (t = 0). (**B**) Uninjected embryo. (**C**) *fry*-MO (15 ng) injected embryo. Note the blastopore formation and subsequent blastopore closure is delayed in *fry*-depleted embryos. Dotted yellow arrows indicate the position and length of the dorsal blastopore lip when it is formed. Dotted white lines in t = 480 min outline the vegetal hemisphere and the blastopore of the embryos. Yellow and blue lines indicate the position of the blastopore from the dorsal and ventral sides, respectively. Note that the blastopore closes eccentric through the ventral side in uninjected embryo and concentric in *fry*-depleted embryo. D: dorsal; V: ventral.

### Fry depletion affects movement of superficial involuting marginal zone cells during gastrulation

Since blastopore closure is driven by multiple processes including convergent thickening, convergent extension, and involution (Keller et al., 2003; Shook et al., 2018), we first investigated the impact of Fry depletion on the motion of the superficial cells during involution (Shook et al., 2004). To achieve this, we injected dorsal blastomeres with *H2B-eGFP* mRNA with or without *fry*-MO and mounted the embryos for *light-sheet* fluorescence microscopy. This technique allowed us to visualize dorsal cell nuclei over several hours with negligible levels of photobleaching in intact, live gastrulating embryos (Movie S2 and Movie S3) (Moretti et al., 2020). We obtained information about the position and motion of hundreds of single cells by performing maximum intensity projections of the raw 3D stacks, which we segmented and used to track nuclei over time (Fig. 4A) (Haubold et al., 2016). In order to measure velocity and persistence of motion of dorsal superficial involuting marginal zone (IMZ) cells, we selected cells that: (i) were tracked for at least 45 minutes, (ii) moved a distance of at least 20 μm and (iii) were closer than 80 μm to the DBL at the last time point that were tracked. We measured the average instantaneous velocity and persistence for the filtered cells as described in the Methods section. First, we evaluated the persistence of each cell as it moved towards the DBL. Persistence defined as the ratio between the linear distance traveled by a cell and the total length of its path, gives a measurement of directionality: cells that move in a trajectory closer to a line will have persistence closer to 1, whereas cells that move in a more erratic trajectory will have lower persistence. We observed that Fry depletion significantly reduced directional persistence with a mean of 0.93 ± 0.06 for cells of embryos control versus a mean of 0.85 ± 0.05 for cells of *fry*-depleted embryos (Fig. 4B). For control embryos, the majority of the cells (> 50%) had persistence values between 1 and 0.95 while very few cells had values below 0.70. By contrast, for *fry*-depleted embryos, less than 20% of the cells fall within the 1 - 0.95 persistence interval while the remaining 80% had lower persistence values. These results indicate that cells trajectories towards the DBL are severely affected in the absence of Fry (Fig. 4C, D). Next, we defined instantaneous velocity as the spatial displacement over time for two consecutive frames. In consistency with the observed blastopore closure delay, Fry depletion significantly reduced the instantaneous velocity of individual cells as they move towards the DBL (Fig. 4E). Next, we thought to investigate whether the spatial organization of cells on the superficial IMZ was affected as a result of the observed velocity and persistence changes. We estimated the distance between nearby cells nuclei as a function of the distance from the DBL. To this end, we used Delaunay triangulation on the nuclei centers to find for each one their closest neighbors and then calculate the mean distance to them (see Methods). Cells were then binned according to the distance from the closest point to the blastopore lip. From this analysis, we found significant differences in the distance to neighbors with dependence on the distance from the blastopore. While cells at distances of 75-95 μm from the DBL showed a similar distance to their neighbor cells in uninjected and *fry*-depleted embryos, cells closer to the DBL (up to 65 μm) presented a significant reduction of the distance to neighbors in *fry*-morphants (Fig. 4F). This result reveals an alteration of the geometrical organization in the pre-involution zone (up to 30 μm from the DBL) of the superficial IMZ as a result of Fry depletion. This observed increase of cell density or “bunching” in the proximity of the involution site in *fry-*depleted embryos (Fig. 4G), could indicate that while superficial IMZ cells are displaying convergence forces, they are not being pulled by inner forces. Together, these results demonstrate a dorsal-requirement for Fry during involution movements and suggest that convergent thickening might be operating in *fry*-morphants allowing for blastopore closure even under conditions where convergent extension is likely defective (Keller et al., 2003; Shook et al., 2018).

**Figure 4.**
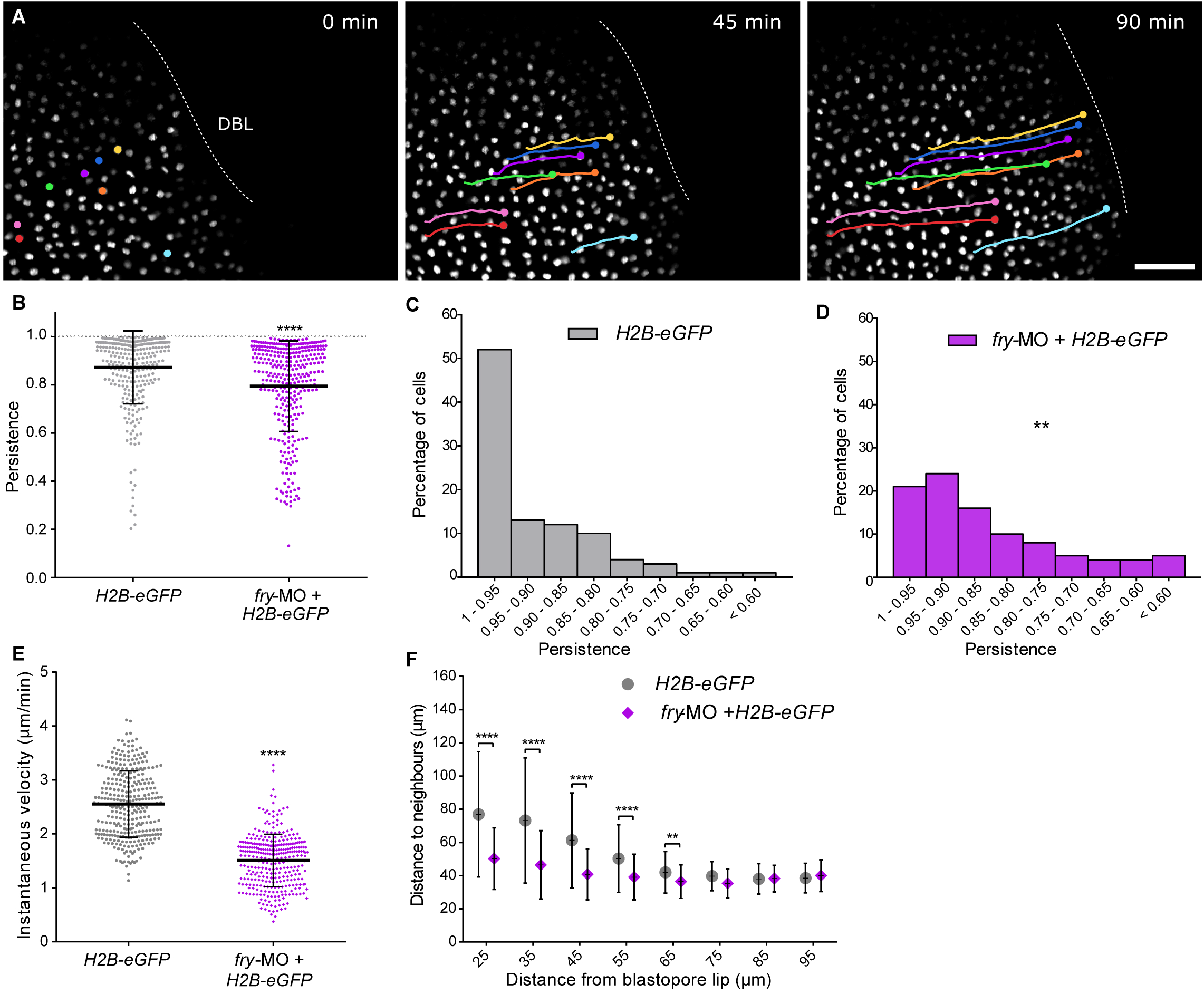
Fry depletion affects the motion of superficial involuting marginal zone cells. (**A**) *Xenopus* 4-cell stage embryos were dorsally injected with *H2B.eGFP* mRNA with or without *fry*-MO (15 ng) and mounted for *light-sheet* fluorescence microscopy at the beginning of gastrulation (St. 10). Time lapse movies were recorded during gastrulation and individual cells were tracked while moving toward the dorsal blastopore lip (DBL) (e.g. nuclei in color. Circles and lines indicate nuclei position and trajectory, respectively). Uninjected embryos (N = 5; total number of tracked cells = 319); *fry*-MO injected embryos (N = 5; total number of tracked cells = 332). N: number of independent experiments. Representative photograms of a time-lapse movie in selected time-points (t) are shown. Scale bar: 100 μm. (**B**) Individual cell persistence measurement was calculated as the ratio between the linear distance traveled by a cell and the total length of its path. Each point represents a single cell. * indicates statistically significant differences between groups, two-tailed Mann Whitney *U*-test (**** *p*<0.0001). The mean and standard deviation are indicated. (**C**) Histogram representing the percentage of cells from control (*H2B.eGFP*) embryos for the different persistence intervals. (**D**) Histogram representing the percentage of cells from *fry-*MO + *H2B.eGFP* embryos for the different persistence intervals. Statistically significant differences were found between groups, *Chi*-square test (** *p*<0.001) (**E**) Individual cell instantaneous velocity measurement. The mean and standard deviation are indicated. Each point represents a single cell. * indicate statistically significant differences between groups, two-tailed Mann Whitney *U*-test (**** *p*<0.0001). (**F**) Average distance from each cell nuclei to the nearest neighbors for all cells within a certain distance region from the blastopore lip (region size = 30 μm; overlapping region size = 5 μm). Distance to neighbors was quantified from the 150 min time point (St. 11.5) of the movies shown in (A). Number of cells in each window was always larger than 100 cells. Data in the graph is presented as means with standard deviation. * indicates statistically significant differences between groups, Kruskal-Wallis test and Dunn’s multiple comparisons test (**** *p*<0.0001, ** *p*<0.01).

### Fry function is necessary for formation of the cleft of Brachet and associated fibronectin network

Before the overt surface movements of gastrulation (Nieuwkoop and Faber, 1994), a deep cleft forms named the cleft of Brachet, separating deep vegetal cells from the more superficial cell layers of the overlying marginal zone (Nieuwkoop and Faber, 1994; Winklbauer and Keller, 1996). In late blastula stages, an ECM rich in fibronectin forms in this cleft (Davidson et al., 2004). At the start of gastrulation, deep layers of the mesendodermal cells move over the base of the cleft, the so-called inner-lip, to initiate involution and move up the fibronectin interface with overlying, e.g. pre-involution cells. There is a distinct behavioral change as cells move over this inner-lip that maintains separation between pre- and post-involution cells. As cells move over this inner lip, they acquire the capacity of remain separated from the pre-involution cells and the ectodermal cells of the blastocoel roof (BCR). This behavioral change can be exposed by tissue separation (Winklbauer and Keller, 1996, Winkbauer 2001).

To assess the cleft of Brachet formation in *fry*-morphants we analyzed hemisected embryos and found that while it forms anteriorly, it is absent near the blastopore lip (Fig. 5A,B). In addition to revealing the presence of the cleft of Brachet, fibronectin fibrils are the main components of the ECM playing an essential role in the regulation of polarized cell protrusions of mesodermal cell (Davidson et al., 2008; Marsden and DeSimone, 2001; Winklbauer and Keller, 1996). In early gastrula *fry*-depleted embryos, we observed that the normally abundant fibronectin fibrils in the cleft were reduced (Fig. 5C-E), while other populations of fibrils on the BCR of *fry-*MO injected embryos were not affected (Fig. 5 C,D). Given that abnormalities in the formation of the cleft of Brachet are frequently associated with tissue separation defects we evaluated tissue separation behavior using BCR assays (Wacker et al., 2000; Winklbauer and Keller, 1996). Involuted mesoderm test aggregates of uninjected and *fry*-MO injected embryos remained on the explanted BCR surface, demonstrating that internalized mesodermal cells are capable of maintaining separation from the explanted BCR cells (Fig. S3). As expected, aggregates isolated from the inner layer of the BCR merged into the explanted BCR surface, as those cells do not express separation behavior (Fig. S3).

**Figure 5.**
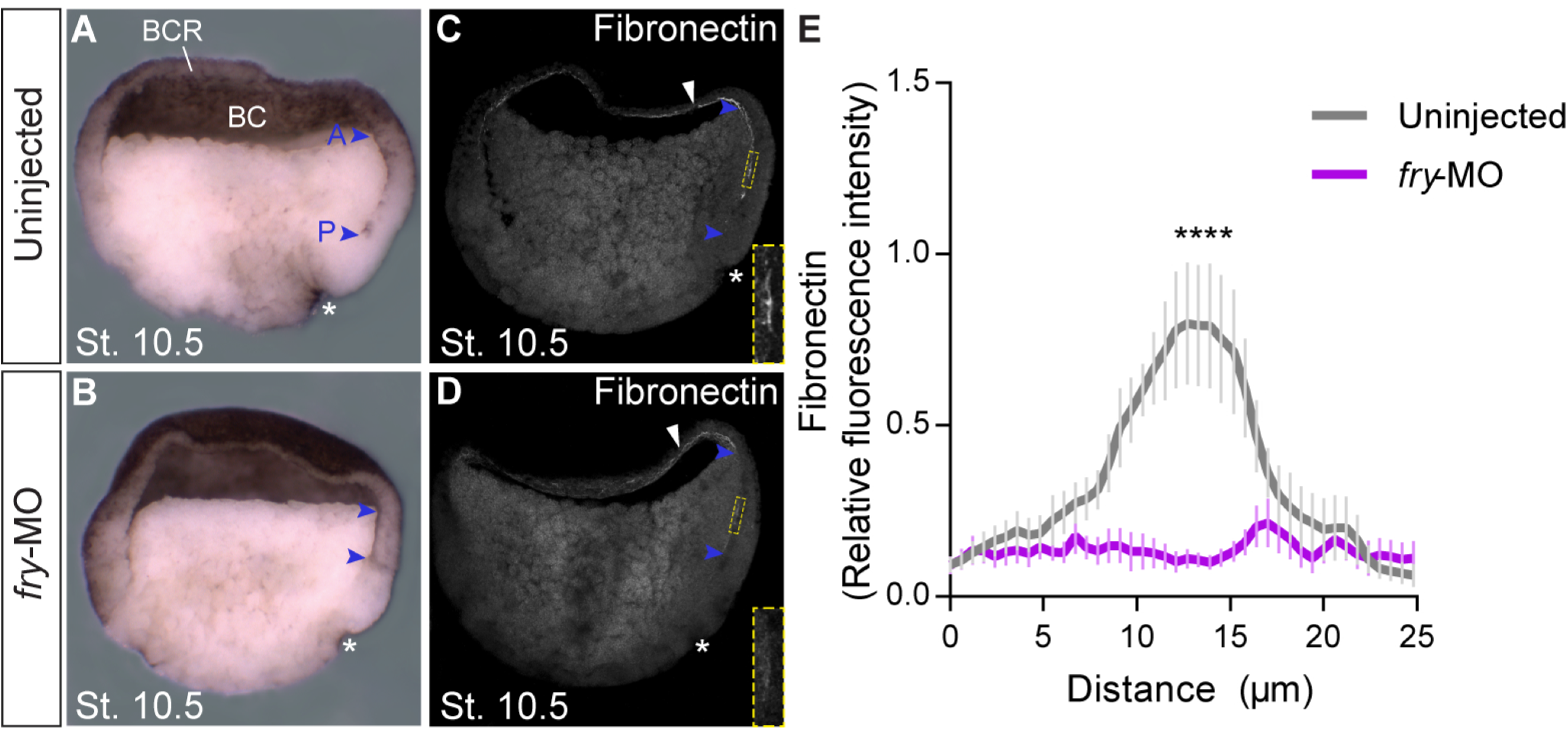
The cleft of Brachet and fibronectin network formation are affected in *fry*-depleted embryos. (**A**, **B**) Early gastrula stage embryos (St. 10.5) hemisections. (**A**) Uninjected embryo (N = 2, n = 26). (**B**) *fry*-MO injected embryo (N = 2; n = 25). * indicate the position of the dorsal blastopore lip; blue arrowheads indicate the anterior (A) and posterior (P) limits of the cleft of Brachet, BC: blastocoel; BCR: blastocoel roof. Representative embryos are shown. (**C**, **D**) Hemisections of fibronectin immunostained early gastrula stage embryos (St. 10.5). (**C**) Uninjected embryo. (**D**) *fry*-MO (15 ng) injected embryo. A rectangle 25 μm width and 100 μm height was drawn at a distance of 300 μm from the dorsal blastopore lip across the cleft of Brachet for fibronectin quantification (dotted lines). A magnification of the quantified area is shown on the right bottom corner. White arrowheads indicate fibronectin presence at the BCR. (**G**) Graph represents the normalized fibronectin abundance across the 25 μm width of the rectangle in uninjected embryos (N = 2; n = 7) and *fry*-MO (15 ng) injected embryos (N = 2; n = 8). Data in the graph are presented as means with standard error. * indicate statistically significant differences between groups, two-tailed Mann Whitney *U*-test (**** *p*<0.0001).

Taken together, our findings show that loss of Fry function does not alter cell affinities measured by the BCR tissue separation assay, however, knocking down Fry impairs the posterior cleft formation and the assembly of the fibronectin fibrils required for attachment and migration of mesodermal cells post-involution.

### Furry is required for dorsal mesoderm convergent extension and cell elongation

Convergent extension within dorsal midline tissues extends the body axis and aids blastopore closure (Shih and Keller, 1992a; Shook et al., 2018). Loss of Fry function inhibits normal blastopore closure (Fig. 3), reduces the levels of fibronectin fibrills in the cleft of Brachet (Fig. 5), which then results in abnormal expression domains of dorsal axial markers (Fig. 2). These observations and the phenotype of *fry*-morphants tailbuds suggest that Fry plays a role in convergent extension of the dorsal mesoderm. To explore this possibility we first sought to quantify convergent extension in intact embryos by measuring the length of *not* expression domain in dorsal midline tissues of whole-embryos at early-gastrula and late-gastrula (Ewald et al., 2004). In untreated control embryos, the expression domain of *not* undergoes antero-posterior elongation and lateral narrowing during these stages (Fig. S4). In accordance with our initial observation (Fig. 2), dorsal mesoderm elongation in late-gastrula was inhibited in the absence of Fry and partially rescued by *FD+LZ* mRNA injection (Fig. S4).

To isolate the process of convergent extension from other processes that also play a role during gastrulation (Shook et al., 2018), we prepared dorsal marginal zone (DMZ) “open-face” explants (Keller et al., 1992). Unlike control explants that elongate as a result of convergent extension (Fig. 6A,B), elongation of DMZ explants prepared from *fry*-depleted embryos was strongly inhibited (Fig. 6C,E). Additionally, co-injection of *FD+LZ* mRNA was able to restore elongation (Fig. 6D,E) consistent with the axis elongation rescue by tailbud stage (Fig. 1) and indicating that defective convergent extension is at least in part, responsible for the shortened axis phenotype.

**Figure 6.**
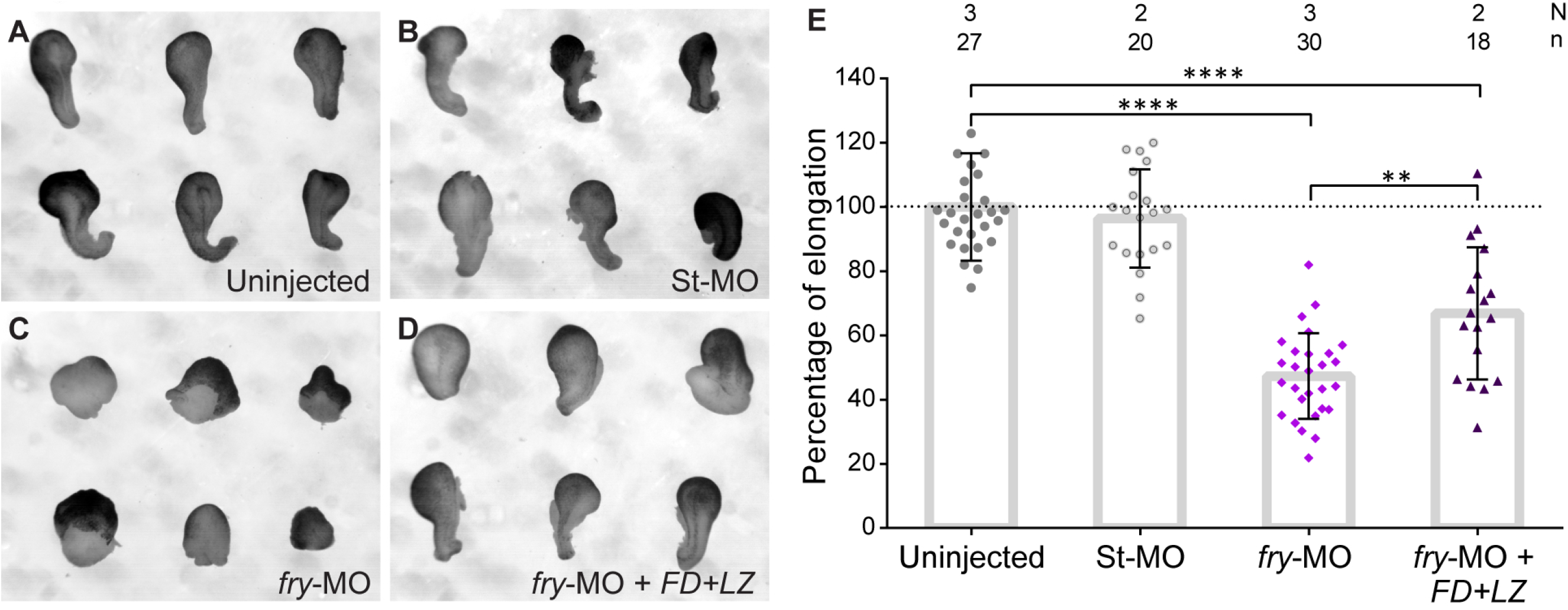
Fry is necessary for convergent extension movements of dorsal mesoderm. (**A**-**D**) Dorsal marginal zone explants were prepared from *Xenopus* embryos at early gastrula stage (St. 10.5) and culture until late neurula stage (St. 19) when the elongation of the explant was evaluated. (**A**) Uninjected embryos. (**B**) Standard Control morpholino (St-MO) (15 ng) injected embryos. (**C**) *fry*-MO (15 ng) injected embryos and (**D**) *fry*-MO (15 ng) + *FD+LZ* mRNA (800 pg) coinjected embryos. Representative explants are shown. (**E**) Percentage of elongation of dorsal marginal zone explants from embryos treated as indicated. Elongation was calculated as the difference between the initial and final length of the explants (St. 10.5 vs. St. 19) relative to the mean of the uninjected group (considered 100% elongation, dotted line). N: number of independent experiments, n: number of embryos. Data in the graph is presented as means with standard deviation. Each point represents a single explant. * indicate statistically significant differences between groups, Kruskal-Wallis test and Dunn’s multiple comparisons test (**** *p*<0.0001, ** *p*<0.01).

To drive convergent extension, mesoderm cells must undergo directed cell rearrangement through mediolateral cell intercalation, a cell behavior marked by mediolateral cell elongation (Shih and Keller, 1992c). To evaluate wherever Fry plays a role in regulating mediolateral cell intercalation, we assessed cell shape in DMZ “open-face” explants isolated from embryos dorsally injected with *fry*-MO and a membrane marker (mem-mScarlet). The degree of cell shape polarization or “polarity index” of dorsal mesoderm cells was measured as the ratio between cell major and minor axes (Fig. 7A-C; Davidson et al., 2006; Feroze et al., 2015). Cells lacking Fry showed a lower polarity index (1.64 ± 0.02) relative to cells from embryos injected with the mRNA alone (1.92 ± 0.03) (Fig. 7C). Interestingly, in the absence of Fry, dorsal mesoderm cells presented with a smaller area than in the control group consistent with the loss of cell elongation (Fig. 7D). These experiments demonstrate that Fry function is required for cell elongation and convergent extension of dorsal mesoderm cells.

**Figure 7.**
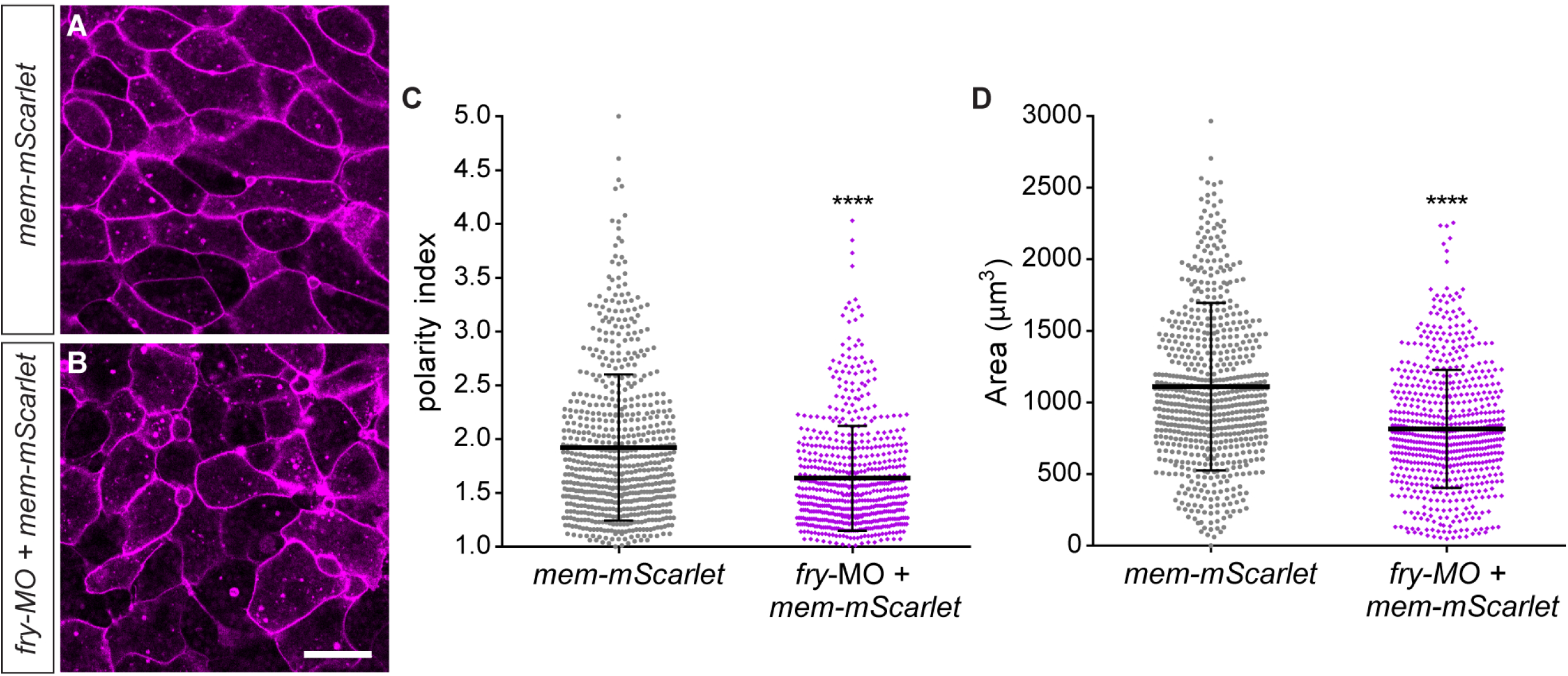
Loss of Fry affects morphological polarity of mesodermal cells. (**A**-**D**) Dorsal marginal zone explants were prepared from *Xenopus* embryos at early gastrula stage (St. 10.5) and imaged at late gastrula stage (St. 13) to assess dorsal mesoderm cell morphology. (**A**) *mem-mScarlet* mRNA injected embryo. (**B**) *fry-*MO (15 ng) + *mem-mScarlet* mRNA coinjected embryo. Scale bar: 100 μm. Representative explants are shown. (**C**) Polarity index measurements of dorsal mesoderm cells calculated as the ratio between cell major axis and minor axis. (**D**) Cellular area measurements of dorsal mesoderm cells. *mem-mScarlet* mRNA injected embryos (N = 2; n = 7; total number of cells analyzed = 660); *fry*-MO + *mem-mScarlet* mRNA coinjected embryos (N = 2; n = 7; total number of cells analyzed = 632). N: number of independent experiments, n: number of embryos. Means and standard deviation are indicated. Each point represents a single cell. * indicate statistically significant differences between groups, two-tailed Mann Whitney *U*-test (**** *p*<0.0001).

### Human NDR1 kinase rescues axis elongation and convergent extension of *fry*-depleted *Xenopus* embryos

In mammalian cells and invertebrates, Fry orthologs genetically and physically interact with NDR kinases (Nagai and Mizuno, 2014). Human NDR1 (hNDR1) kinase activation is regulated by phosphorylation of two sites (Ser281 and Thr444) and through its association with mps one binder (MOB) proteins and Fry proteins (Chiba et al., 2009; Devroe et al., 2004; Xiong et al., 2018). However, there is no evidence of either physical or functional interactions between Fry and NDR1 in vertebrate development. First, we evaluated the axis development in embryos injected with an mRNA encoding a hyperactive form of human NDR1 named *hNDR1-PIF* which mimics an active kinase, and another of a kinase-dead version named *hNDR1.kd* (Cook et al., 2014). Similar to the phenotype of *fry*-morphants, both *hNDR1* mRNAs caused axis defects in ~50% of the embryos characterized by reduction of head structures and mild shortened axis phenotype (Fig. 8C,D,G). The observed phenotypes demonstrate that dorsal overexpression of human NDR1 functional variants affect axis development in *Xenopus*.

**Figure 8.**
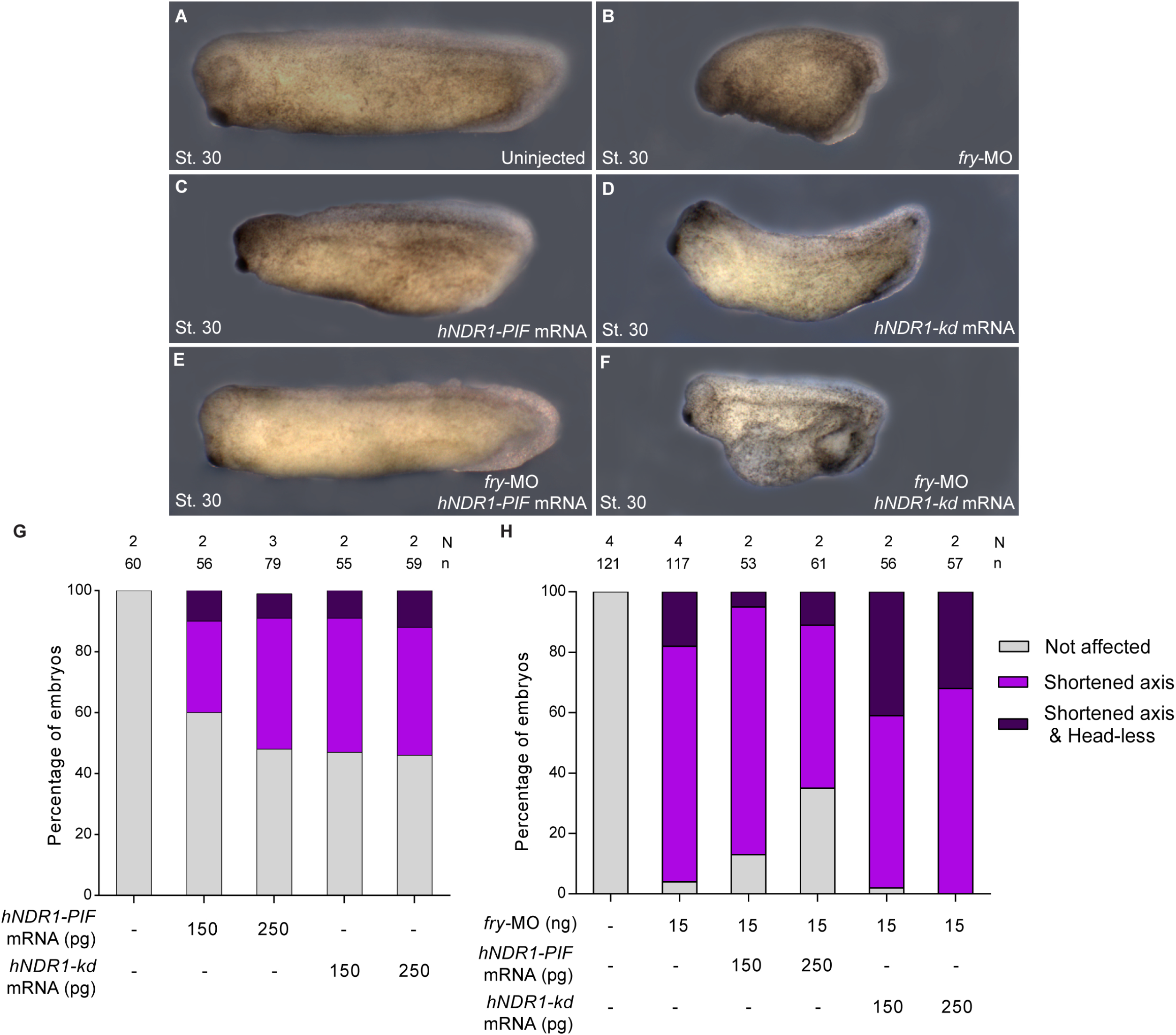
hNDR1 rescues *fry*-depletion axis phenotype. (**A**-**F**) 4-cell *Xenopus* embryos were injected into both dorsal blastomeres as indicated and fixed at St. 30. (**A**) Uninjected embryo. (**B**) *fry*-MO (15 ng) injected (**C**) *hNDR1-PIF* mRNA (250 pg) injected embryo. (**D**) *hNDR1-kd* mRNA (250 pg) injected embryo. (**E**) *fry*-MO (15 ng) + *hNDR1-PIF* mRNA (250 pg) coinjected embryo. (**F**) *fry*-MO (15 ng) + *hNDR1-kd* mRNA (250 pg) coinjected embryo. Representative embryos are shown. (**G, H**) Quantitation of the percentage of embryos showing the different phenotypes: “Not affected”, “Shortened axis” or “Shortened axis & Head-less” phenotypes. Data on graph is presented as mean. N: number of independent experiments, n: number of embryos.

Next we evaluated the functional interaction of hNDR1 and Fry in axis development by performing rescue experiments coinjecting *fry-*MO with *hNDR1-PIF* mRNA. The tailbud *fry*-morphant phenotype was partially rescued by *hNDR1-PIF* in a dose-dependent manner (Fig. 8E,H). However, the expression of the variant *hNDR1.kd* failed to rescue the *fry*-depletion phenotype (Fig. 8F,H), demonstrating that NDR1 kinase activity is required for the rescue. Additionally, a higher percentage of embryos showed a more severe phenotype when coinjected with *fry*-MO and *hNDR1.kd* arguing in favor of a functional interaction (Fig. 8H).

We went further and evaluated whether hNDR1 was able to rescue convergent extension in *fry*-depleted embryos. Analysis of DMZ explants elongation showed that *hNDR1-PIF* partially compensates for Fry loss-of-function (Fig. 9). However, *hNDR1-kd* coinjected explants elongated subtly, but the differences in the percentage of elongation with the *fry*-MO group was not significant (Fig. 9D,E) indicating that hyperactive hNDR1 and not the kinase-dead hNDR1 is able to compensate for absence of Fry in the recovery of convergent extension. Our results are in agreement with previous studies showing that Fry is critical for the activation of NDR1 kinase (Chiba et al., 2009; Nagai and Mizuno, 2014) and demonstrate that the axis elongation and convergent extension phenotypes in *fry*-morphants can be explained at least in part, by NDR1 kinase function.

**Figure 9.**
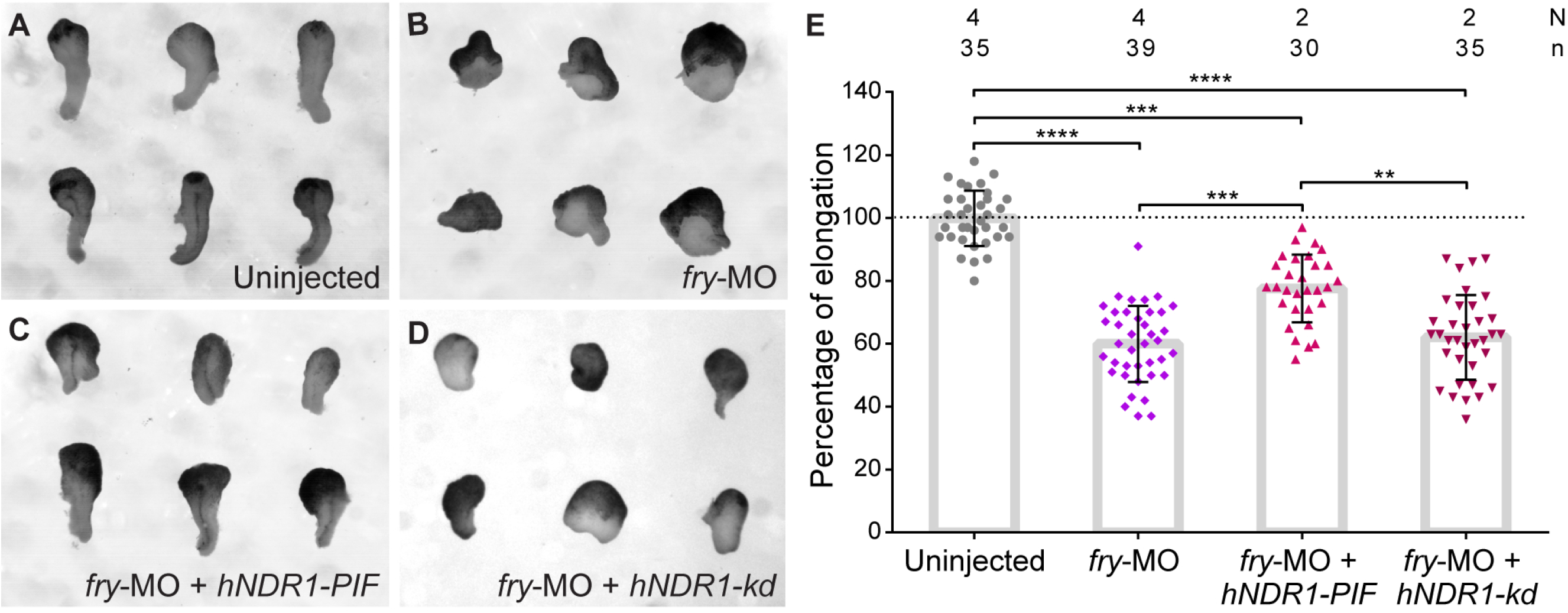
Hyperactive hNDR1 rescues convergent extension after Fry loss-of-function. (**A**-**D**) Dorsal marginal zone explants were prepared from *Xenopus* embryos at early gastrula stage (St. 10.5) and culture until late neurula stage (St. 19). (**A**) Uninjected embryos, (**B**) *fry*-MO (15 ng) injected embryos, (**C**) *fry*-MO (15 ng) + *hNDR1-PIF* mRNA (250 pg) coinjected embryos and (**D**) *fry*-MO (15 ng) + *hNDR1-kd* mRNA (250 pg) coinjected embryos. Representative explants are shown. (**E**) Percentage of elongation of dorsal marginal zone explants from embryos treated as indicated. Elongation was calculated as indicated on Fig. 6. Data in the graph is presented as means with standard deviation. Each point represents a single explant. * indicate statistically significant differences between groups, Kruskal-Wallis test and Dunn’s multiple comparisons test (**** *p*<0.0001). N: number of independent experiments, n: number of embryos.

## Discussion

### Fry transcripts and protein cellular localization

In *Xenopus*, *fry* has been described as a maternally expressed gene (Goto et al., 2010). When analyzing its distribution in the blastula as well as earlier stages and unfertilized eggs (Fig. S1A, A’ and our unpublished observation), we observed an enrichment of the transcripts in the animal half (Fig. S1). Understanding the functional relevance of this asymmetric distribution will require further studies and could provide insight into the mechanisms that regulate the coupling of asymmetry in oogenesis with the establishment of asymmetry in the embryo. At the beginning of gastrulation, *fry* transcripts are present in the dorsal and ventral marginal zones supporting a role in the initial gastrulation movements (Fig. S1). Consistently, we found that Fry loss-of-function affects the movements and spatial configuration of the superficial IMZ cells (Fig. 4). As gastrulation proceeds, *fry* mRNA is enriched in the axial and paraxial mesoderm and the deep ectodermal layer, all tissues that undergo convergent extension (Fig. S1). Fry transcripts are also present in the ventral marginal zone of the early gastrula however, understanding its function in ventral tissues will require further investigation. In this work, we confirmed that *fry*-depleted embryos present reduced anterior-head structures and shortened axis (Goto et al., 2010). Our studies demonstrate that while mesoderm induction is not impaired and trunk structures such as somites and notochord are formed (Figs 2,S2) Fry is required for the morphogenetic behavior of midline tissues (Figs S4,6).

Similarly to endogenous Fry protein localization in *Drosophila* pupal wings cells (He et al., 2005), follicular cells of the egg chamber (Horne-Badovinac et al., 2012) and in HeLa cells (Chiba et al., 2009), we show that a Fry-GFP fusion protein is present in the cytoplasm and cell membrane of dorsal mesodermal cells of mid-gastrula *Xenopus* embryos (Fig. S1). In a previous report, Fry-GFP was also detected in the nucleus of isolated mesodermal *Xenopus* cells (Goto et al., 2010), however in the explants analyzed we did not detect nuclear protein. Fry protein found in the nuclei of *Drosophila* salivary gland and fat body cells (He et al., 2005), argues in favor of Fry being a highly mobile protein which cellular localization might be cell type-specific and time-specific. Here, our results point to a cytoplasmic function of Fry in dorsal mesoderm cells during gastrulation.

### Fry and the regulation of gastrulation movements in *Xenopus*

Blastopore closure in the amphibian embryo is achieved as the result of large scale tissue morphogenesis shaped by mechanical forces (Shook et al., 2018). There are two intrinsic convergence behaviors of the IMZ that contribute to blastopore closure: convergent thickening and convergent extension. While convergent extension is exclusively conducted by the presumptive dorsal tissues of the mid-gastrula, convergent thickening operates in all preinvoluting circumblastoporal tissues in a symmetric fashion (Keller et al., 1985; Shook et al., 2018). As it was demonstrated in UV ventralized embryos, convergent thickening can drive blastopore closure on its own in the absence of convergent extension nevertheless less efficiently (Scharf and Gerhart, 1980; Shook et al., 2018). We observed that, similar to ventralized embryos, dorsal Fry depletion alters the initial blastopore formation, blastopore closure dynamics, and blastopore closing position in these embryos, suggesting that convergent thickening operates even in the absence of Fry (Fig. 3, Movies S1, S2 and S3). Additionally, our cell tracking recordings of *fry*-depleted embryos reveal alterations in the movement of superficial IMZ cells and the spatial organization of the cells near the blastopore lip (Fig. 4), revealing a lack of coordination between the forces that operate in the system. Together, our findings show for the first time a role of Fry as a regulator of cell movements during gastrulation in vertebrates.

### Fry and the intercellular and cell-ECM interactions during gastrulation

Prior to gastrulation, vegetal endoderm rotates toward the dorsal ectoderm (vegetal rotation) placing the mesoderm next to the ectoderm. Cells from both germ layers display tissue separation behavior and the cleft of Brachet is established (Wacker et al., 2000; Winklbauer and Schürfeld, 1999). As mesoderm cells involute, they express the separation behavior and are able to migrate along the ectodermal BCR without mixing. Thus, the anterior portion of the cleft of Brachet is the result of vegetal rotation and the posterior part is formed during mesoderm involution (Wacker et al., 2000). We observed that the posterior part of the cleft of Brachet is not visible in *fry*-depleted embryos (Fig. 5B) despite mesoderm cells lacking Fry having the capacity to remain separate from the BCR cells (Fig. S3). In this regard, the fibronectin fibrillar matrix normally assembled at the cleft of Brachet of the early gastrula is missing in theses embryos, despite its presence at the BCR (Fig. 5). Fibronectin ECM plays a critical role in the regulation and maintenance of gastrulation movements since fibronectin knock-down results in delayed blastopore closure (Davidson et al., 2006). Fibronectin fibrillary matrix at the cleft of Brachet provides physical support for mesendoderm cells migration (Davidson et al., 2004; Davidson et al., 2008; Rozario et al., 2009), as well as a source of signaling either through its own cellular receptor integrin or as surface for growth factors strategical accumulation (Davidson et al., 2002; Plouhinec et al., 2013). The abnormal development of anterior structures in *fry*-depleted embryos might be the result of alterations in leading-edge mesodermal cells migratory kinetics due to physical changes in the fibronectin matrix. The establishment and maintenance of mediolaterally directed protrusive activity that drives mediolateral cell intercalation of dorsal mesoderm cells undergoing convergent extension requires integrin-fibronectin recognition (Davidson et al., 2006). As in fibronectin knock-down, convergent extension is disrupted in *fry*-depleted embryos (Fig. 6E). However, DMZ cells lacking Fry failed to elongate in an exogenously supplied fibronectin substrate (Fig. 7) suggesting the existence of an additional mechanism affecting the morphogenetic behavior of these cells.

### Fry and the Wnt/PCP signaling pathway

Convergent extension is the result of polarized cell movements characterized by the mediolateral intercalation of cells along the axis, resulting in elongation of the embryo. The non-canonical Wnt/PCP signaling pathway is key to convergent extension as most of its molecules and downstream effectors contribute to polarized cell behaviors and their dysregulation results in convergent extension failure and defective blastopore closure (Butler and Wallingford, 2017; Darken et al., 2002; Goto and Keller, 2002; Goto et al., 2005; Keller, 2002; Medina et al., 2000; Moon et al., 1993; Park and Moon, 2002; Sokol, 1996; Takeuchi et al., 2003; Wallingford et al., 2000) however other pathways and molecules are involved in these processes too (Chung et al., 2007; Luxardi et al., 2010; Ulmer et al., 2017). In addition to regulating cell polarity and the coordination of morphogenetic behaviors, PCP complex regulates polarized ECM deposition required for convergent extension (Goto et al., 2005; Gray et al., 2011). Our experiments show that convergent extension is impaired in *fry*-depleted embryos and both elongation and mediolateral alignment of the cells engaged in convergent extension are lost. Moreover, we also observed a reduction of fibronectin fibrillar matrix assembly along the mesoderm surface prior to convergent extension, which resembles PCP pathway perturbations (Figs 6, 7) (Goto et al., 2005). Likewise, the similarity we observed between the phenotypes associated with Fry loss-of-function and gain-of-function by overexpression of the short chimeric version FD+LZ is also common for core PCP components, which might suggest both pathways having common downstream effectors and having a role in the same cellular processes (Djiane et al., 2000; Goto and Keller, 2002; Goto et al., 2005; He and Adler, 2002; Keller, 2002; Kraft et al., 2012; Tada and Smith, 2000; Takeuchi et al., 2003; Wallingford, 2012; Wallingford et al., 2000; Winklbauer et al., 2001). While we have not looked in this study at effector proteins of the PCP pathway such as the small GTPases of the Rho family, in *Drosophila* sensory neurons, genetic interaction with Rho proteins and PCP components have been shown for the Fry/tricornered (*trc*, ortholog of NDR1 in *Drosophila*) pathway regulating dendritic self-avoidance and tiling (Emoto et al., 2004; Matsubara et al., 2011). Further studies will be required to determine the level of physical and functional interaction between Fry and PCP pathways components in *Xenopus*.

### Fry and NDR1 functional interaction

Multiple studies in invertebrates and ours here in *Xenopus*, point to a role of Fry protein in cellular mechanisms driving morphogenesis. Fry was originally identified in *Drosophila melanogaster* where its mutation causes disorganized epidermal cell morphology (Cong et al., 2001). A very similar phenotype found in *tricornered* gene mutants suggested that these proteins physically interact and function in a common pathway (Cong et al., 2001; Du and Novick, 2002; He et al., 2005). Despite the fact that genetic studies in yeast, nematode and fruit fly have revealed a critical role of Fry in the control of cell morphogenesis and cell polarity associated with the activation of NDR kinases, the molecular and cellular mechanisms are not well understood. Our rescue experiments demonstrate that human NDR1 can rescue the *fry*-morphants phenotype, indicating that hyperactive hNDR1 can partially compensate for the loss of Fry function in *Xenopus* (Figs 8, 9). The specific function of *Xenopus* NDR1 (Stk38) in axis elongation and convergent extension movements in *Xenopus* will require further investigation however, our results argue in favor of a conserved functional interaction between Fry and NDR kinases in animal development.

Several questions related to the cellular and molecular mechanisms by which Fry regulates morphogenetic movements during gastrulation remain unanswered. However, based on our results and previously published data, we can formulate three hypothesis to explain the observed morphogenetic defects yet not exclusive. Based on the migratory defects and loss of cell polarization, one possibility is that Fry regulates the cytoskeleton dynamics. In *Drosophila*, Fry and Trc have been found to alter actin organization during egg chamber elongation and hair wing morphogenesis in addition to regulate microtubule sliding in neurons (Horne-Badovinac et al., 2012; Norkett et al., 2020). Another possibility is that Fry regulates cell adhesion of *Xenopus* mesodermal cells to the ECM and the assembly of the ECM itself affecting their migratory behavior. In this sense, Fry promotes dendrite attachment to the ECM in *Drosophila* da neurons, however the mechanism remains elusive (Han et al., 2012). Whether in our system Fry is required for intercellular adhesion and/or cell-ECM adhesion via integrins remains to be determined. A third possibility we have considered, is that Fry regulates the orientation of cell divisions, as precisely oriented divisions are important to maintain cell polarity, polarized cell movement, axis elongation and several other developmental processes (Concha and Adams, 1998; Gong et al., 2004; Kieserman and Wallingford, 2009; Théry and Bornens, 2006). This last hypothesis is based on Fry’s reported function in chromosome alignment and mitotic spindle orientation in mammalian cells (Chiba et al., 2009; Ikeda et al., 2012; Nagai et al., 2013). Likewise the identification of upstream and downstream factors will provide important information about Fry’s mechanism of action.

## Materials and methods

### Ethics Statement

This study was carried out in strict accordance with the recommendations in the Guide for the Care and Use of Laboratory Animals of the NIH. The protocol was approved by the Institutional Animals Care and Use Committee (IACUC) of the School of Applied and Natural Sciences, University of Buenos Aires, Argentina (Protocol #64).

### *Xenopus* embryos preparation

*Xenopus laevis* embryos were obtained by natural mating. Adult frog reproductive behavior was induced by injection of human chorionic gonadotropin hormone. Eggs were collected, de-jellied in 3% cysteine (pH 8.0), maintained in 0.1 X Marc’s Modified Ringer’s (MMR) solution and staged according to Nieuwkoop and Faber (Nieuwkoop and Faber, 1994). The embryos were placed in 3% ficoll prepared in 1 X MMR for microinjection.

### Constructs for mRNA synthesis

Human NDR1 constructs were generously provided by Alexander Hergovich. The hNDR1-PIF (constitutive hyperactive) and hNDR1-kd (kinase-dead) (Cook et al., 2014) cDNAs were excised from pcDNA3.HA.hNDR1-PIF and pcDNA3.HA.hNDR1-kd (K118A) by *BamH*I and *Xho*I digestion and cloned into pCS2+. Fry-GFP construct was generously provided by Toshiyasu Goto (Goto et al., 2010). The constructs for pCS2+.H2B-eGFP (Gong et al., 2004) and pCS2+.HA.FD+LZ (Espiritu et al., 2018) had been previously described. pCS2+.mem-mScarlet was generated using mScarlet (Bindels et al., 2017) cloned into pCS2+ with a membrane-targeting domain (mem) corresponding to the farnesylation motif from human HRas.

### Morpholino and mRNA microinjections

Capped mRNAs for *fry-GFP*, *mem-mScarlet, H2B-eGFP*, *HA.FD+LZ*, *hNDR1-PIF* and *hNDR1-kd* were *in vitro* transcribed using a mMessage mMachine kit (Ambion) following linearization with *Not*I. Fry morpholino *(fry*-MO) (Gene Tools, LLC) sequence and specificity have been previously published (Goto et al., 2010). A Morpholino Standard Control oligo (St-MO) was used as a negative control (Gene Tools, LLC). Morpholinos (MO) or mRNAs were injected into both dorsal blastomeres of 4-cell embryos targeting the DMZ. *Fry*-MO was injected at 5 - 15 ng per embryo and St-MO at 15 ng per embryo. The doses of injected mRNAs per embryo were as follows: *fry-GFP* (1.2 ng), *mem-mScarlet* (125 pg), *HA.FD+LZ* (200 - 800 pg), *H2B-eGFP* (500 pg), *hNDR1-PIF* and *hNDR1-kd* (150 - 250 pg).

### *In situ* hybridization and immunostaining

Whole-mount *in situ* hybridization was carried out as previously described (Gawantka et al., 1998). *Fry* (Dharmacon), *chordin* (gift from Edward De Robertis) and the *myoD* constructs (gift from Oliver Wessely) were linearized as previously described (Cirio et al., 2011; Espiritu et al., 2018; Sasai et al., 1994). *Brachyury* construct (gift from Neil Hukriede) was linearized with *Bgl*II and *xnot* construct (gift from David Kimelman) was linearized with *Hind*III. All linearized constructs were transcribed with T7 for antisense probe synthesis. For whole-mount immunostaining with MZ15 (DSHB Cat# MZ15, RRID:AB_760352) and 12/101 (DSHB Cat# 12/101, RRID:AB_531892) we followed the protocol previously described (Cirio et al., 2011). Fibronectin immunostaining with 4H2 (DSHB Cat# 4H2, RRID:AB_2721949) and subsequent clearing and mounting for confocal microscopy was performed as previously described (Davidson et al., 2004). Hemisections were cut before staining and imaged with an Olympus FV confocal microscope. Fibronectin was measured with ImageJ software (https://fiji.sc/). A 25 μm width and 100 μm high rectangle was drawn around the cleft of Brachet 300 μm away from the DBL. Fluorescence intensity was quantified across the width and relativized to the mean intensity in order to compare between independent experiments. For preparation of histological slides, embryos processed for *in situ* hybridization or immunostaining were post-fixed in Bouin’s solution, dehydrated, cleared in xylene, embedded in paraffin and sectioned at 15 μm (Espiritu et al., 2018). Xenbase (http://www.xenbase.org/, RRID:SCR_003280) was used as source of information on gene expression, developmental stages and anatomy.

### Image analysis

Fixed whole embryos were photographed externally with a Leica DFC420 camera attached to a Leica L2 stereoscope. Histological slides were imaged using a digital camera (Infinity 1; Lumera Corporation) attached to a light-field microscope (CX31: Olympus,). For gastrulation time-lapse movies, late blastula (St. 9) uninjected and *fry-MO* injected embryos were selected and transferred to custom acrylic chamber with 0.1 X MMR. Gastrulation was recorded for approximately 13 hours (18°C). Images were taken every 3 minutes using an Imaging Source SMK 23G445 camera attached to a Zeiss Axiovert S 100 scope. Gastrulation measurements were quantified in fixed embryos using ImageJ software (https://fiji.sc/). Blastopore closure was calculated as the ratio of the blastopore area over the area of the vegetal hemisphere of the embryo. Elongation of dorsal midline tissues evaluated as the length *not* expression domain over the length of the whole-embryo was determined as previously described (Ewald et al., 2004).

### Embryos microsurgery (DMZ explants)

Following microinjection, early gastrula embryos (St. 10.5) were transferred to 1 X Modified Barth’s Saline (MBS) solution. Vitelline membranes were removed with forceps and DMZ explants were microsurgically isolated using hair tools (Feroze et al., 2015; Shawky et al., 2018). Explants were immobilized with cover glass and cultured until co-cultured intact siblings reached late neurula stage (St. 19). The percentage of elongation was calculated as previously described (Wallingford et al., 2000). For confocal microscopy, explants were prepared in Danilchik’s for Amy (DFA) culture media and mounted in customs acrylic chambers prepared in advance with 20 μg/ml fibronectin (Sigma) (Zhou et al., 2010). The explants were imaged with a Leica Microsystems SP5 confocal microscope. For Fry-GFP visualization in mesodermal cells, DMZ explants were allowed to heal for 30-40 minutes before imaging. For cell polarity measurement, explants were cultured until sibling embryos reached late gastrula stage (St. 13/14). Polarity index and cell area were calculated as previously described (Feroze et al., 2015) using ImageJ software (https://fiji.sc/).

### Light-sheet fluorescence microscopy

Following microinjection, late blastula embryos (St. 9) were placed in pre-warmed (22°C) 0.2 % agarose (Biodynamics) prepared in 0.1 X MMR. Embryos were drawn into 4 cm length FEP tubes (Rotilabo, D-76185 inner diameter 1.58 mm, outer diameter 3.18 mm) with a needle and syringe (Kaufmann et al., 2012). The tube was plugged with solid 1% agarose and the samples were imaged using a custom-built light-sheet fluorescence microscope (LSFM) (Moretti et al., 2020). This microscope uses a cylindrical lens to generate a static Gaussian light-sheet that is focused into a sample using an illumination objective (Olympus UMPLFLN 10XW, 0.3 NA). The resulting light-sheet has a thickness of around 5 μm over a field of view of around 1 mm x 1 mm. The emitted fluorescence is collected using a detection objective (Olympus UMPLFLN 20XW, 0.5 NA) followed by a filter wheel, a tube lens and an sCMOS camera (Andor Zyla). Samples were scanned across the light-sheet using a stepper motor and the whole microscope was controlled using a custom-made graphical interface. 3D stacks of injected embryos were acquired upon visualization of the blastopore formation and every 3 minutes using an axial step of 5 μm over a distance of 400 - 600 μm. The resulting images were segmented and tracked using *ilastik* (Haubold et al., 2016). We analyzed the tracking results using custom-made Python scripts. We defined a cell’s instantaneous velocity *v* as *v* = Δ*r*/Δt, where Δr is the cell’s spatial displacement over two consecutive frames and Δt the corresponding frame rate (3 minutes in all our experiments). We defined persistence of motion as the ratio between the linear distance traveled by a cell and the total length of its migration path (Matthews et al., 2008). Thus, persistence gives a measurement of motion directionality: cells that move in a trajectory closer to a line will have persistence closer to 1, whereas cells that move in a more erratic trajectory will have lower persistence. The distance to the nearest neighbors was calculated as follows. For a single time-point (t = 150 min), the center of each nuclei as reported by *ilastik* was fed into the Delaunay algorithm (Python 3.6, SciPy 1.3) to obtain the list of nearest neighbors (NN). The euclidean distance from each nuclei to their NN was calculated and then averaged. The distance of each cell from the blastopore lip was defined as the closest distance from the nuclei to a curve that was manually drawn over the lip for the frame of interest (region size = 30 μm; overlapping region size = 5 μm).

### BCR assays

Following microinjection, early gastrula embryos (St. 10.5) were transferred to 1 X Modified Barth’s Saline (MBS) solution. BCR assays to assess tissue separation behavior were performed as previously described (Wacker et al., 2000). Briefly, prechordal or deep layer ectodermal cells derived from uninjected or *fry*-MO injected embryos were dissected into small cell aggregates (test aggregates) and placed upon an uninjected BCR. Explants were immobilized with cover glass and separation behavior was scored after 45 min.

### Statistical Analysis

Numbers of embryos (n) and independent experimental replicates (N) for animal studies are stated in graphs. Statistical analysis for Figures 4B, 4E, 5, S3 and 7 was performed using two-tailed Mann Whitney *U*-test (**** *p*<0.0001). Statistical analysis for Figures 3, 4F, S4, 6 and 9 were performed using Kruskal-Wallis test and means between groups were compared using Dunn’s multiple comparisons test (**** *p*<0.0001). Statistical analysis for Figure 4C,D were performed using *Chi*-square test (** *p*<0.001). For all statistical analyses we used Prism6, GraphPad Software, Inc.

## Acknowledgments

We would like to thank Lirane Moutinho and Hernán Diego Martin for technical support, Neil A. Hukriede, Silvia L. Lopez, Daniel Hochbaum and Aitana M. Castro Colabianchi for reagents and discussions, Alexander Hergovich for the hNDR1-PIF and hNDR1-kd plasmids.

## Competing interests statement

The authors declare no competing financial interests.

## Funding

MCC laboratory was supported by the Agencia Nacional de Promoción Científica y Tecnológica of Argentina (PICT-2013-0381), the Concejo Nacional de Investigaciones Científicas y Técnicas (CONICET) of Argentina (PIP-11220150100577CO). HEG laboratory was supported by Fondo para la Investigación Científica y Tecnológica (2013-1301,2014-3658), Max-Planck-Gesellschaft (Partner Group) and Secretaría de Ciencia y Técnica (20020170100755BA). ASC and BM were supported by the CONICET Doctoral Fellowship Program. LAD and CS were supported by the National Institutes of Health (NICHD R01 HD044750 and NHLBI R01 R01 HL136566).

## References

Bindels, D. S., Haarbosch, L., van Weeren, L., Postma, M., Wiese, K. E., Mastop, M., Aumonier, S., Gotthard, G., Royant, A., Hink, M. A., et al. (2017). mScarlet: a bright monomeric red fluorescent protein for cellular imaging. Nat Methods 14, 53–56.

Butler, M. T. and Wallingford, J. B. (2017). Planar cell polarity in development and disease. Nat Rev Mol Cell Biol 18, 375–388.

Chiba, S., Ikeda, M., Katsunuma, K., Ohashi, K. and Mizuno, K. (2009). MST2- and Furry-mediated activation of NDR1 kinase is critical for precise alignment of mitotic chromosomes. Curr Biol 19, 675–681.

Chung, H. A., Yamamoto, T. S. and Ueno, N. (2007). ANR5, an FGF Target Gene Product, Regulates Gastrulation in Xenopus. Curr. Biol.

Cirio, M. C., Hui, Z., Haldin, C. E., Cosentino, C. C., Stuckenholz, C., Chen, X., Hong, S. K., Dawid, I. B. and Hukriede, N. A. (2011). Lhx1 is required for specification of the renal progenitor cell field. PLoS One 6, e18858.

Concha, M. L. and Adams, R. J. (1998). Oriented cell divisions and cellular morphogenesis in the zebrafish gastrula and neurula: A time-lapse analysis. Development.

Cong, J., Geng, W., He, B., Liu, J., Charlton, J. and Adler, P. N. (2001). The furry gene of Drosophila is important for maintaining the integrity of cellular extensions during morphogenesis. Development 128, 2793–2802.

Cook, D., Hoa, L. Y., Gomez, V., Gomez, M. and Hergovich, A. (2014). Constitutively active NDR1-PIF kinase functions independent of MST1 and hMOB1 signalling. Cell Signal 26, 1657–1667.

Darken, R. S., Scola, A. M., Rakeman, A. S., Das, G., Mlodzik, M. and Wilson, P. A. (2002). The planar polarity gene strabismus regulates convergent extension movements in Xenopus. EMBO J 21, 976–985.

Davidson, L. A., Hoffstrom, B. G., Keller, R. and DeSimone, D. W. (2002). Mesendoderm extension and mantle closure in Xenopus laevis gastrulation: Combined roles for integrin α5β1, fibronectin, and tissue geometry. Dev. Biol.

Davidson, L. A., Keller, R. and DeSimone, D. W. (2004). Assembly and remodeling of the fibrillar fibronectin extracellular matrix during gastrulation and neurulation in Xenopus laevis. Dev Dyn 231, 888–895.

Davidson, L. A., Marsden, M., Keller, R. and Desimone, D. W. (2006). Integrin alpha5beta1 and fibronectin regulate polarized cell protrusions required for Xenopus convergence and extension. Curr Biol 16, 833–844.

Davidson, L. A., Dzamba, B. D., Keller, R. and Desimone, D. W. (2008). Live imaging of cell protrusive activity, and extracellular matrix assembly and remodeling during morphogenesis in the frog, Xenopus laevis. Dev. Dyn. 237, 2684–2692.

Devroe, E., Erdjument-Bromage, H., Tempst, P. and Silver, P. A. (2004). Human Mob proteins regulate the NDR1 and NDR2 serine-threonine kinases. J. Biol. Chem.

Djiane, A., Riou, J. F., Umbhauer, M., Boucaut, J. C. and Shi, D. L. (2000). Role of frizzled 7 in the regulation of convergent extension movements during gastrulation in Xenopus laevis. Development.

Du, L. L. and Novick, P. (2002). Pag1p, a novel protein associated with protein kinase Cbk1p, is required for cell morphogenesis and proliferation in Saccharomyces cerevisiae. Mol Biol Cell 13, 503–514.

Emoto, K., He, Y., Ye, B., Grueber, W. B., Adler, P. N., Jan, L. Y. and Jan, Y. N. (2004). Control of dendritic branching and tiling by the Tricornered-kinase/Furry signaling pathway in Drosophila sensory neurons. Cell 119, 245–256.

Espiritu, E. B., Crunk, A. E., Bais, A., Hochbaum, D., Cervino, A. S., Phua, Y. L., Butterworth, M. B., Goto, T., Ho, J., Hukriede, N. A., et al. (2018). The Lhx1-Ldb1 complex interacts with Furry to regulate microRNA expression during pronephric kidney development. Sci Rep 8, 16029.

Ewald, A. J., Peyrot, S. M., Tyszka, J. M., Fraser, S. E. and Wallingford, J. B. (2004). Regional requirements for Dishevelled signaling during Xenopus gastrulation: separable effects on blastopore closure, mesendoderm internalization and archenteron formation. Development 131, 6195–6209.

Feroze, R., Shawky, J. H., von Dassow, M. and Davidson, L. A. (2015). Mechanics of blastopore closure during amphibian gastrulation. Dev Biol 398, 57–67.

Gallegos, M. E. and Bargmann, C. I. (2004). Mechanosensory neurite termination and tiling depend on SAX-2 and the SAX-1 kinase. Neuron 44, 239–249.

Gawantka, V., Pollet, N., Delius, H., Vingron, M., Pfister, R., Nitsch, R., Blumenstock, C. and Niehrs, C. (1998). Gene expression screening in Xenopus identifies molecular pathways, predicts gene function and provides a global view of embryonic patterning. Mech. Dev.

Gong, Y., Mo, C. and Fraser, S. E. (2004). Planar cell polarity signalling controls cell division orientation during zebrafish gastrulation. Nature.

Goto, T. and Keller, R. (2002). The planar cell polarity gene Strabismus regulates convergence and extension and neural fold closure in Xenopus. Dev. Biol.

Goto, T., Davidson, L., Asashima, M. and Keller, R. (2005). Planar cell polarity genes regulate polarized extracellular matrix deposition during frog gastrulation. Curr Biol 15, 787–793.

Goto, T., Fukui, A., Shibuya, H., Keller, R. and Asashima, M. (2010). Xenopus furry contributes to release of microRNA gene silencing. Proc Natl Acad Sci U S A 107, 19344–19349.

Gray, R. S., Roszko, I. and Solnica-Krezel, L. (2011). Planar Cell Polarity: Coordinating Morphogenetic Cell Behaviors with Embryonic Polarity. Dev. Cell.

Han, C., Wang, D., Soba, P., Zhu, S., Lin, X., Jan, L. Y. and Jan, Y. N. (2012). Integrins Regulate Repulsion-Mediated Dendritic Patterning of Drosophila Sensory Neurons by Restricting Dendrites in a 2D Space. Neuron.

Haubold, C., Schiegg, M., Kreshuk, A., Berg, S., Koethe, U. and Hamprecht, F. A. (2016). Segmenting and tracking multiple dividing targets using ilastik. Adv. Anat. Embryol. Cell Biol.

He, B. and Adler, P. N. (2002). The genetic control of arista lateral morphogenesis in Drosophila. Dev Genes Evol 212, 218–229.

He, Y., Fang, X., Emoto, K., Jan, Y. N. and Adler, P. N. (2005). The tricornered Ser/Thr protein kinase is regulated by phosphorylation and interacts with furry during Drosophila wing hair development. Mol Biol Cell 16, 689–700.

Hergovich, A., Stegert, M. R., Schmitz, D. and Hemmings, B. A. (2006). NDR kinases regulate essential cell processes from yeast to humans. Nat Rev Mol Cell Biol 7, 253–264.

Hirata, D., Kishimoto, N., Suda, M., Sogabe, Y., Nakagawa, S., Yoshida, Y., Sakai, K., Mizunuma, M., Miyakawa, T., Ishiguro, J., et al. (2002). Fission yeast Mor2/Cps12, a protein similar to Drosophila furry, is essential for cell morphogenesis and its mutation induces Wee1-dependent G2 delay. EMBO J. 21, 4863–4874.

Horne-Badovinac, S., Hill, J., Gerlach 2nd, G., Menegas, W. and Bilder, D. (2012). A screen for round egg mutants in Drosophila identifies tricornered, furry, and misshapen as regulators of egg chamber elongation. G3 2, 371–378.

Ikeda, M., Chiba, S., Ohashi, K. and Mizuno, K. (2012). Furry protein promotes aurora A-mediated Polo-like kinase 1 activation. J Biol Chem 287, 27670–27681.

Kaufmann, A., Mickoleit, M., Weber, M. and Huisken, J. (2012). Multilayer mounting enables long-term imaging of zebrafish development in a light sheet microscope. Development 139, 3242–3247.

Keller, R. (2002). Shaping the vertebrate body plan by polarized embryonic cell movements. Science (80-.). 298, 1950–1954.

Keller, R. and Shook, D. (2008). Dynamic determinations: patterning the cell behaviours that close the amphibian blastopore. Philos Trans R Soc L. B Biol Sci 363, 1317–1332.

Keller, R. E., Danilchik, M., Gimlich, R. and Shih, J. (1985). The function and mechanism of convergent extension during gastrulation of Xenopus laevis. J Embryol Exp Morphol 89 Suppl, 185–209.

Keller, R., Shih, J. and Domingo, C. (1992). The patterning and functioning of protrusive activity during convergence and extension of the Xenopus organiser. Dev Suppl 81–91.

Keller, R., Davidson, L., Edlund, A., Elul, T., Ezin, M., Shook, D. and Skoglund, P. (2000). Mechanisms of convergence and extension by cell intercalation. Philos Trans R Soc L. B Biol Sci 355, 897–922.

Keller, R., Davidson, L. A. and Shook, D. R. (2003). How we are shaped: the biomechanics of gastrulation. Differentiation 71, 171–205.

Kieserman, E. K. and Wallingford, J. B. (2009). In vivo imaging reveals a role for Cdc42 in spindle positioning and planar orientation of cell divisions during vertebrate neural tube closure. J. Cell Sci.

Kraft, B., Berger, C. D., Wallkamm, V., Steinbeisser, H. and Wedlich, D. (2012). Wnt-11 and Fz7 reduce cell adhesion in convergent extension by sequestration of PAPC and C-cadherin. J Cell Biol 198, 695–709.

Leptin, M. (2005). Gastrulation movements: the logic and the nuts and bolts. Dev Cell 8, 305–320.

Luxardi, G., Marchal, L., Thome, V. and Kodjabachian, L. (2010). Distinct Xenopus Nodal ligands sequentially induce mesendoderm and control gastrulation movements in parallel to the Wnt/PCP pathway. Development 137, 417–426.

Marsden, M. and DeSimone, D. W. (2001). Regulation of cell polarity, radial intercalation and epiboly in Xenopus: Novel roles for integrin and fibronectin. Development 128, 3635–3647.

Matsubara, D., Horiuchi, S. Y., Shimono, K., Usui, T. and Uemura, T. (2011). The seven-pass transmembrane cadherin Flamingo controls dendritic self-avoidance via its binding to a LIM domain protein, Espinas, in Drosophila sensory neurons. Genes Dev.

Matthews, H. K., Marchant, L., Carmona-Fontaine, C., Kuriyama, S., Larraín, J., Holt, M. R., Parsons, M. and Mayor, R. (2008). Directional migration of neural crest cells in vivo is regulated by Syndecan-4/Rac1 and non-canonical Wnt signaling/RhoA. Development 135, 1771–1780.

Medina, A., Reintsch, W. and Steinbeisser, H. (2000). Xenopus frizzled 7 can act in canonical and non-canonical Wnt signaling pathways: Implications on early patterning and morphogenesis. Mech. Dev.

Moon, R. T., Campbell, R. M., Christian, J. L., McGrew, L. L., Shih, J. and Fraser, S. (1993). Xwnt-5A: A maternal Wnt that affects morphogenetic movements after overexpression in embryos of Xenopus laevis. Development.

Moretti, B., Müller, N. P., Wappner, M. and Grecco, H. E. (2020). Compact and reflective light-sheet microscopy for long-term imaging of living embryos. Appl. Opt. 59, D89–D94.

Nagai, T. and Mizuno, K. (2014). Multifaceted roles of Furry proteins in invertebrates and vertebrates. J Biochem 155, 137–146.

Nagai, T., Ikeda, M., Chiba, S., Kanno, S. and Mizuno, K. (2013). Furry promotes acetylation of microtubules in the mitotic spindle by inhibition of SIRT2 tubulin deacetylase. J Cell Sci 126, 4369–4380.

Nelson, B., Kurischko, C., Horecka, J., Mody, M., Nair, P., Pratt, L., Zougman, A., McBroom, L. D. B., Hughes, T. R., Boone, C., et al. (2003). RAM: A conserved signaling network that regulates Ace2p transcriptional activity and polarized morphogenesis. Mol. Biol. Cell.

Nieuwkoop, P. D. and Faber, J. (1994). Normal table of Xenopus Laevis. New York: Garland Publishing.

Norkett, R., del Castillo, U., Lu, W. and Gelfand, V. I. (2020). Ser/Thr kinase Trc controls neurite outgrowth in Drosophila by modulating microtubule-microtubule sliding. Elife.

Park, M. and Moon, R. T. (2002). The planar cell-polarity gene stbm regulates cell behaviour and cell fate in vertebrate embryos. Nat. Cell Biol.

Plouhinec, J. L., Zakin, L., Moriyama, Y. and De Robertis, E. M. (2013). Chordin forms a self-organizing morphogen gradient in the extracellular space between ectoderm and mesoderm in the Xenopus embryo. Proc. Natl. Acad. Sci. U. S. A.

Rozario, T., Dzamba, B., Weber, G. F., Davidson, L. A. and DeSimone, D. W. (2009). The physical state of fibronectin matrix differentially regulates morphogenetic movements in vivo. Dev. Biol.

Sasai, Y., Lu, B., Steinbeisser, H., Geissert, D., Gont, L. K. and De Robertis, E. M. (1994). Xenopus chordin: a novel dorsalizing factor activated by organizer-specific homeobox genes. Cell 79, 779–790.

Scharf, S. R. and Gerhart, J. C. (1980). Determination of the dorsal-ventral axis in eggs of Xenopus laevis: Complete rescue of uv-impaired eggs by oblique orientation before first cleavage. Dev. Biol.

Shawky, J. H., Balakrishnan, U. L., Stuckenholz, C. and Davidson, L. A. (2018). Multiscale analysis of architecture, cell size and the cell cortex reveals cortical f-actin density and composition are major contributors to mechanical properties during convergent extension. Dev.

Shih, J. and Keller, R. (1992a). Patterns of cell motility in the organizer and dorsal mesoderm of Xenopus laevis. Development 116, 915–930.

Shih, J. and Keller, R. (1992b). The epithelium of the dorsal marginal zone of Xenopus has organizer properties. Development 116, 887–899.

Shih, J. and Keller, R. (1992c). Cell motility driving mediolateral intercalation in explants of Xenopus laevis. Development 116, 901–914.

Shook, D. R., Majer, C. and Keller, R. (2004). Pattern and morphogenesis of presumptive superficial mesoderm in two closely related species, Xenopus laevis and Xenopus tropicalis. Dev. Biol. 270, 163–185.

Shook, D. R., Kasprowicz, E. M., Davidson, L. A. and Keller, R. (2018). Large, long range tensile forces drive convergence during Xenopus blastopore closure and body axis elongation. Elife 7,.

Smith, J. C., Price, B. M. J., Green, J. B. A., Weigel, D. and Herrmann, B. G. (1991). Expression of a xenopus homolog of Brachyury (T) is an immediate-early response to mesoderm induction. Cell.

Sokol, S. Y. (1996). Analysis of Dishevelled signalling pathways during Xenopus development. Curr Biol 6, 1456–1467.

Tada, M. and Smith, J. C. (2000). Xwnt11 is a target of Xenopus Brachyury: regulation of gastrulation movements via Dishevelled, but not through the canonical Wnt pathway. Development 127, 2227–2238.

Takeuchi, M., Nakabayashi, J., Sakaguchi, T., Yamamoto, T. S., Takahashi, H., Takeda, H. and Ueno, N. (2003). The prickle-related gene in vertebrates is essential for gastrulation cell movements. Curr Biol 13, 674–679.

Théry, M. and Bornens, M. (2006). Cell shape and cell division. Curr. Opin. Cell Biol.

Ulmer, B., Tingler, M., Kurz, S., Maerker, M., Andre, P., Monch, D., Campione, M., Deissler, K., Lewandoski, M., Thumberger, T., et al. (2017). A novel role of the organizer gene Goosecoid as an inhibitor of Wnt/PCP-mediated convergent extension in Xenopus and mouse. Sci Rep 7, 43010.

Von Dassow, G., Schmidt, J. E. and Kimelman, D. (1993). Induction of the Xenopus organizer: Expression and regulation of Xnot, a novel FGF and activin-regulated homeo box gene. Genes Dev. 7, 355–366.

Wacker, S., Grimm, K., Joos, T. and Winklbauer, R. (2000). Development and control of tissue separation at gastrulation in Xenopus. Dev Biol 224, 428–439.

Wallingford, J. B. (2012). Planar cell polarity and the developmental control of cell behavior in vertebrate embryos. Annu Rev Cell Dev Biol 28, 627–653.

Wallingford, J. B., Rowning, B. A., Vogeli, K. M., Rothbacher, U., Fraser, S. E. and Harland, R. M. (2000). Dishevelled controls cell polarity during Xenopus gastrulation. Nature 405, 81–85.

Wallingford, J. B., Fraser, S. E. and Harland, R. M. (2002). Convergent extension: the molecular control of polarized cell movement during embryonic development. Dev Cell 2, 695–706.

Winklbauer, R. and Keller, R. E. (1996). Fibronectin, mesoderm migration, and gastrulation in Xenopus. Dev Biol 177, 413–426.

Winklbauer, R. and Schürfeld, M. (1999). Vegetal rotation, a new gastrulation movement involved in the internalization of the mesoderm and endoderm in Xenopus. Development.

Winklbauer, R., Medina, A., Swain, R. K. and Steinbeisser, H. (2001). Frizzled-7 signalling controls tissue separation during Xenopus gastrulation. Nature.

Xiong, S., Lorenzen, K., Couzens, A. L., Templeton, C. M., Rajendran, D., Mao, D. Y. L., Juang, Y. C., Chiovitti, D., Kurinov, I., Guettler, S., et al. (2018). Structural Basis for Auto-Inhibition of the NDR1 Kinase Domain by an Atypically Long Activation Segment. Structure.

Zallen, J. A., Peckol, E. L., Tobin, D. M. and Bargmann, C. I. (2000). Neuronal cell shape and neurite initiation are regulated by the Ndr kinase SAX-1, a member of the Orb6/COT-1/warts serine/threonine kinase family. Mol Biol Cell 11, 3177–3190.

Zhou, J., Kim, H. Y., Wang, J. H. and Davidson, L. A. (2010). Macroscopic stiffening of embryonic tissues via microtubules, RhoGEF and the assembly of contractile bundles of actomyosin. Development 137, 2785–2794.

